# Comparative and population genomics approaches reveal the basis of adaptation to deserts in a small rodent

**DOI:** 10.1101/856310

**Authors:** Anna Tigano, Jocelyn P. Colella, Matthew D. MacManes

## Abstract

Organisms that live in deserts offer the opportunity to investigate how species adapt to environmental conditions that are lethal to most plants and animals. In the hot deserts of North America, high temperatures and lack of water are conspicuous challenges for organisms living there. The cactus mouse (*Peromyscus eremicus*) displays several adaptations to these conditions, including low metabolic rate, heat tolerance, and the ability to maintain homeostasis under extreme dehydration. To investigate the genomic basis of desert adaptation in cactus mice, we built a chromosome-level genome assembly and resequenced 26 additional cactus mouse genomes from two locations in southern California (USA). Using these data, we integrated comparative, population, and functional genomic approaches. We identified 16 gene families exhibiting significant contractions or expansions in the cactus mouse compared to 17 other Myodontine rodent genomes, and found 232 sites across the genome associated with selective sweeps. Functional annotations of candidate gene families and selective sweeps revealed a pervasive signature of selection at genes involved in the synthesis and degradation of proteins, consistent with the evolution of cellular mechanisms to cope with protein denaturation caused by thermal and hyperosmotic stress. Other strong candidate genes included receptors for bitter taste, suggesting a dietary shift towards chemically defended desert plants and insects, and a growth factor involved in lipid metabolism, potentially involved in prevention of dehydration. Understanding how species adapted to the recent emergence of deserts in North America will provide an important foundation for predicting future evolutionary responses to increasing temperatures, droughts and desertification in the cactus mouse and other species.

## Introduction

For decades, researchers have been intrigued by adaptation, or the process by which organisms become better fitted to their environments. To this end, scientists have devoted substantial efforts to this issue and have successfully elucidated how natural selection has shaped organismal phenotypes in response to environmental pressures (Berner and Salzburger, 2015; Cooke et al., 2013; Linnen et al., 2009; Nachman et al., 2003; Savolainen et al., 2013). Given their influence on metabolism, water availability and ambient temperature are environmental factors relevant to all organisms and are also of growing concern within the context of anthropogenically-induced global climate change and increasing desertification (IPCC, 2018). Studying how animals that are currently living in hot and dry environments have adapted to those conditions is one approach for helping to predict the potential impacts of increasing temperatures and aridity (Hoelzel, 2010; Somero, 2010).

Despite the challenging conditions, a wide variety of organisms have evolved adaptations to live in hot deserts. These adaptations include changes in behavior to avoid dehydration, excessive solar radiation, and heat (e.g., nocturnal life and sheltering in burrows) and a suite of anatomical modifications to dissipate heat (e.g., long body parts and pale colors). Some of the most striking adaptations are at the physiological level and help to either minimize water loss through efficient excretion and reabsorption of water (Schmidt-Nielsen, 1964; Schmidt-Nielsen and Schmidt-Nielsen, 1952) or compensate for lack of environmental water via enhanced production of metabolic water from nutrient oxidation (Takei et al., 2012; Walsberg, 2000). While these adaptations to desert life have been described in several species (for a review on small mammals see Walsberg, 2000) and are important under current climate predictions (IPCC, 2018), the genetic underpinnings of these traits are less well known.

Genomic studies on camels (*Camelus bactrianus*) have provided substantial evidence related to the genomic basis of adaptation to deserts. For example, analysis of the camel genome showed an enrichment of fast-evolving genes involved in lipid and carbohydrate metabolism, potentially linked to energy production and storage in a food-scarce environment (Bactrian Camels Genome Sequencing and Analysis Consortium et al., 2012; Wu et al., 2014). Transcriptome analysis of renal cortex and medulla in control and water-restricted camels showed a strong response to dehydration in genes involved in water reabsorption and glucose metabolism (Wu et al., 2014). Overall, genes in the arachidonic acid pathway seem to play a role in desert adaptation in both camels and desert sheep (Bactrian Camels Genome Sequencing and Analysis Consortium et al., 2012; Yang et al., 2016). This pathway regulates water retention and reabsorption in the kidney, primarily through changes in reno-vascular tone. Aquaporins, transmembrane water channel proteins, are also involved in water reabsorption and urine concentration, and changes in their expression levels have been associated with dry environments in kangaroo rats (Marra et al., 2014, 2012) and the Patagonian olive mouse (Giorello et al., 2018).

The cactus mouse (*Peromyscus eremicus*) is native to the deserts of southwestern North America and displays a suite of adaptations to this extreme environment. Cactus mice have behavioral and anatomical adaptations for heat avoidance and dissipation, such as a nocturnal lifestyle, larger ears, and aestivation (Macmillen, 1965). They have also evolved lower metabolic rates, which result in a reduction in water loss, and resistance to heat stress compared to other generalist *Peromyscus* spp. (Murie, 1961). For example, while several desert rodents produce concentrated urine (Schmidt-Nielsen, 1964; Schmidt-Nielsen and Schmidt-Nielsen, 1952), the cactus mouse is essentially anuric (Kordonowy et al., 2017), which indicates its extreme efficiency of renal water reabsorption. Kordonowy et al. (2017) showed through experimental manipulation of water availability that captive cactus mice were behaviorally and physiologically intact after three days of severe acute dehydration. Gene expression profiling of kidneys highlighted a starvation-like response at the cellular level in dehydrated mice, despite access to food, and strong differential expression of *Cyp4* genes, which are part of the arachidonic acid metabolism pathway (MacManes, 2017). Although these results indicate some degree of convergent evolution with other desert-adapted mammals (Bactrian Camels Genome Sequencing and Analysis Consortium et al., 2012; Takei et al., 2012; Yang et al., 2016), they are limited to expressed genes under particular experimental conditions and in one tissue type only.

Deserts in southwest North America formed relatively recently, only after the retreat of the Pleistocene ice sheets that covered most of the continent during the Last Glacial Maximum approximately 10,000 years ago (Pavlik, 2008). Because the ability to detect genomic signatures of selection depends on coalescent time and effective population size (Nielsen et al., 2005), and given the recent emergence of North American deserts, the footprint of these recent adaptations should continue to be evident in contemporary cactus mouse genomes. Whole genome analyses allow us to detect signatures of selection associated with life in the desert across the complete set of cactus mouse genes, regardless of expression patterns. Further, they allow for analysis of intergenic areas, and for characterization of genomic features that may promote or hinder adaptive evolution, such as the distribution of standing genetic variation and repetitive elements.

To identify genes associated with desert adaptation and to investigate the factors affecting adaptation using the cactus mouse as a model, we first generated a chromosome-level genome assembly and then integrated comparative, population, and functional genomics approaches. As dehydration is a primary challenge desert animals face, we expected to identify signatures of selection associated with metabolism and sodium-water balance (i.e.adaptations that either enhance production of metabolic water or prevent fluid loss via excretion) in line with previous studies in the cactus mouse and other desert-adapted species (Bactrian Camels Genome Sequencing and Analysis Consortium et al., 2012; Giorello et al., 2018; Marra et al., 2014; Takei et al., 2012; Wu et al., 2014; Yang et al., 2016). Our analyses of gene family evolution and selective sweeps point instead to regulation of protein synthesis and degradation as the main target of selection. While we find strong support for an evolutionary response in perception of bitter taste and lipid metabolism, we do not identify an extensive signal of selection at genes linked to water-sodium balance at the whole-genome level.

## Results

### Genome assembly and annotation

Illumina and PacBio reads yielded a draft genome assembly of 2.7 Gb and an N50 of 1.3 Mb. Scaffolding with Hi-C data increased contiguity 100-fold and yielded 24 chromosome-sized scaffolds for a total assembly size of 2.5 Gb, plus 173 Mb of unplaced scaffolds. The final assembly contained 92.9% complete BUSCOs, with 1.2% of the genes duplicated and 3.7% missing. Together these statistics indicate that the cactus mouse genome assembly is highly contiguous, complete and non-redundant (Table 2).

**Table 1.**
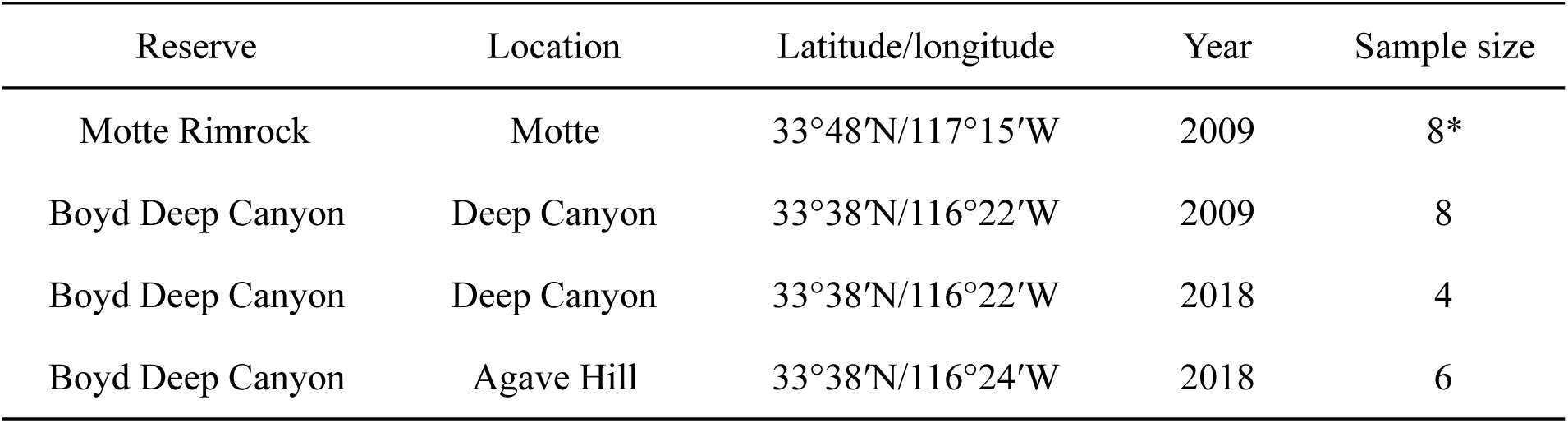
Details on sampling locations of individuals sequenced for population genomics analyses. * denotes that the individual outlier was sampled here, and effective sample size was reduced to 7 after outlier removal

| Reserve | Location | Latitude/longitude | Year | Sample size |
| --- | --- | --- | --- | --- |
| Motte Rimrock | Motte | 33°48'N/117°15'W | 2009 | 8* |
| Boyd Deep Canyon | Deep Canyon | 33°38'N/116°22'W | 2009 | 8 |
| Boyd Deep Canyon | Deep Canyon | 33°38'N/116°22'W | 2018 | 4 |
| Boyd Deep Canyon | Agave Hill | 33°38'N/116°24'W | 2018 | 6 |

**Table 2.** Summary of assembly statistics before and after scaffolding with Hi-C data.

|  | Pre Hi-C assembly | Post Hi-C assembly |
| --- | --- | --- |
| Number of scaffolds | 7650 | 24 (+ 6,785 unplaced scaffolds) |
| Total size of scaffolds | 2.7 Gb | 2.5 Gb (+ 173 Mb unplaced sequence) |
| Longest scaffold | 13.7 Mb | 203.4 Mb |
| Scaffold N50/L50 | 1.3 Mb/530 | 120 Mb/9 |
| Gaps % | 5.51 | 3.53 |

Whole genome alignment to the *P. maniculatus* genome revealed the presence of several intrachromosomal differences between the two species, but no large inversions, translocations, or interchromosomal rearrangements were evident at the resolution granted by *mummer4* (Supplementary Figure 1). This, in combination with a conserved number of chromosomes supported by both karyotype characterization (Smalec et al., 2019) and genome assemblies, indicates that genome structure is highly conserved between these *Peromyscus* species.

We annotated 18,111 protein-coding genes. *Repeatmasker* masked 35% of the genome as repetitive. LINE1 and LTR elements alone constituted 21% of the repeats. Total proportion of repeats and relative proportion of different repeats classes were similar across eight *Peromyscus* species (Supplementary Table 2).

### Genomic differentiation across space and time

Our PCA showed that the first two principal components (PCs) explained 13.32% of the variation present across 26 cactus mice. Both the PCA (Supplementary Figure 2) and the MDS (Figure 1A) clearly separated individuals from Motte and Deep Canyon Reserve. Within the Deep Canyon Reserve, no differentiation at the temporal or microspatial scale was observed (Supplementary Figure 2, Figure 1A).

**Figure 1.**
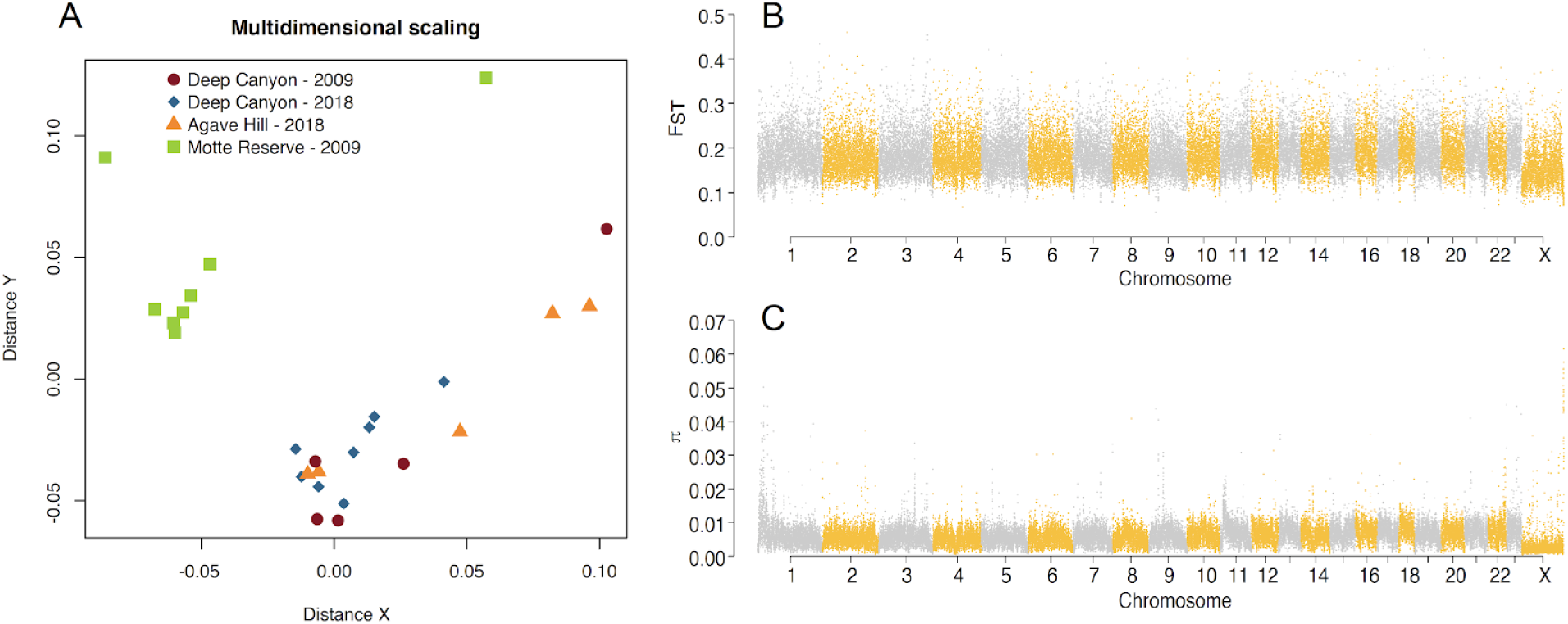
Diversity and differentiation in the cactus mouse. a) MDS plot showing relative distance among individuals based on downsampling to a single base at 43.7 million variable sites. Note outlier from Motte on the far left side of the plot. b) Manhattan plot showing patterns of differentiation based on F_ST_ between Motte and Deep Canyon Reserves (after outlier removal). c) Manhattan plot showing patterns of nucleotide diversity π from all samples combined (after outlier removal).

One individual from Motte appeared distinct from other individuals included in the analysis. As we could not ascertain the reason for such behavior, e.g., technical artifact, taxonomic misidentification, hybridization, etc., this individual was excluded from further analyses.

Differentiation between Motte and Deep Canyon Reserve populations was high, with an average F_ST_ value of 0.19 and 0.14 across the 23 autosomes and the X chromosome, respectively. F_ST_ calculated in 50 kb windows ranged from 0.06 to 0.46 with 95% of the windows ranging from 0.12 and 0.28 (Figure 1B).

### Sequence and structural standing genetic variation

A total of 1,875,915,109 variant and invariant sites, representing 75% of the genome, were included in our analyses. We identified 43,695,428 SNPs with high-confidence (one every 43 bp – 2.3% of all sites). Global genome-wide π was 6×10^-3^. π was lowest on the X chromosome and seemed to increase from chromosome 1 to 23 (Figure 1C). In fact, chromosome length was a strong negative predictor of nucleotide diversity at each autosome (R^2^ = 0.65, F = 39.58, p < 0.001; Figure 2).

**Figure 2.**
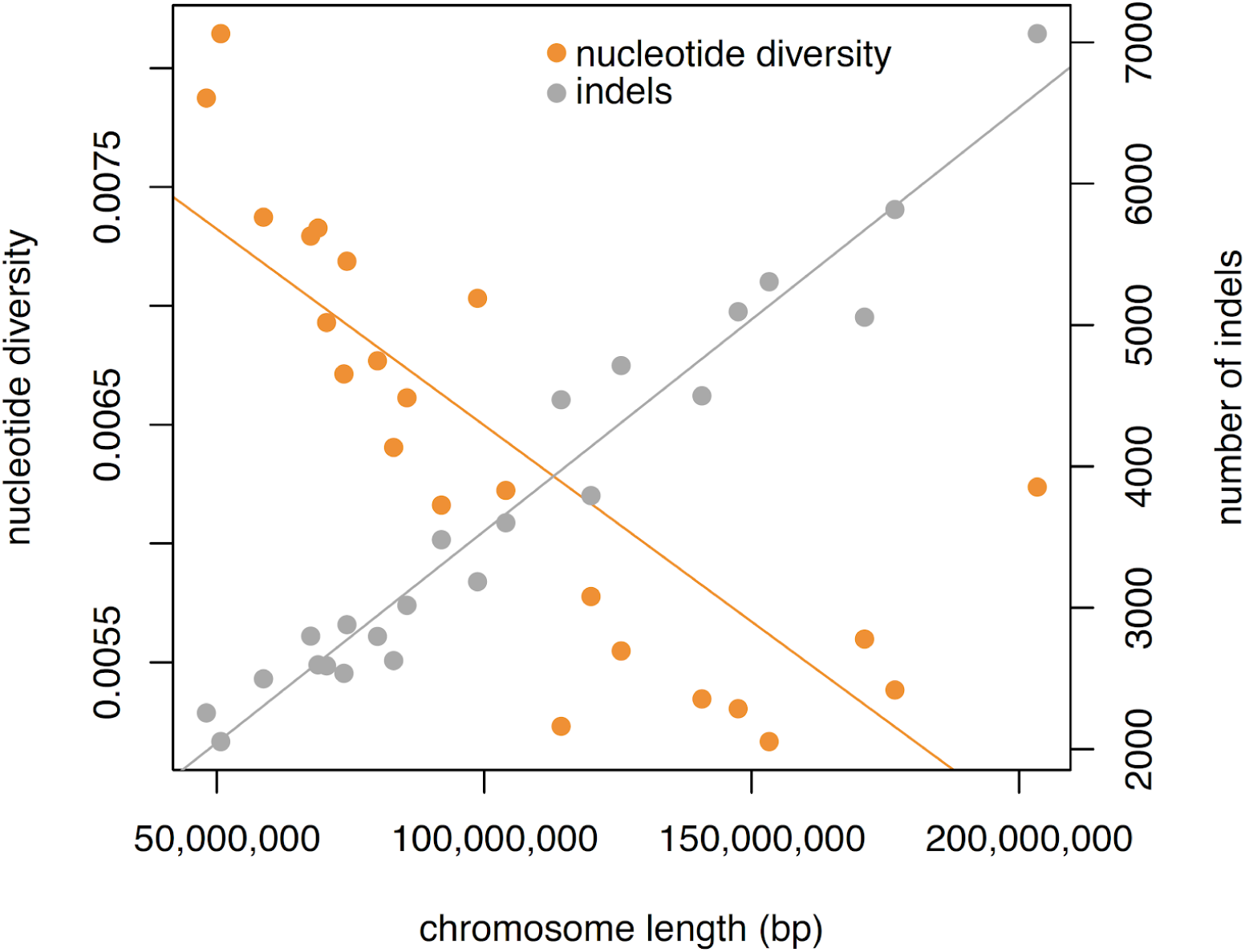
Plot showing mean nucleotide diversity and number of indels as a function of chromosome length (p < 0.001 in both cases, albeit with opposite trends).

A large area of elevated nucleotide diversity (∼17 Mb long) was evident at the beginning of chromosome 1. Conserved synteny with other *Peromyscus* species and unequivocal support from the Hi-C contact map strongly indicated that the assembly was correct for chromosome 1. To begin to understand the genome-level processes that may have generated this pattern, we calculated depth of coverage in chromosomes 1 and 2 - a reference chromosome that did not show similar large regions of elevated π - using a subset of the shotgun data and the number and proportion of repetitive elements in 50 kb windows (Supplementary Figure 3). Depth of sequencing in this unusual area of chromosome 1 was higher than in the rest of chromosome 1 (up to 7x higher) and compared to chromosome 2 (10% higher overall), and showed a similar peak as the one shown for nucleotide diversity (Supplementary Figure 3). The number and proportion of repetitive elements were both higher in this area relative to other parts of chromosome 1, and all of chromosome 2 (Supplementary Figure 3). Together, these analyses suggest that this area is highly repetitive, rather than containing a misassembled large duplication.

Analysis of the 10x Genomics data using *LongRanger* resulted in an estimated mean DNA molecule length of only 13,620 bp (ideal is > 40,000 bp), and the number of linked reads per molecule was six, much lower than the ideal threshold of 13. As short molecules can negatively impact the detection of large structural variants and generate many false positives, we adopted a conservative approach by reporting only short indels (41-29,527 bp). We identified a total of 87,640 indels between the reference and an individual from the same population. Indels affected 101 Mb of the total genome assembly, which represents 4% of the total sequence. Number of indels per chromosome was strongly positively correlated with chromosome size (R^2^ = 0.95, F = 438.7, p < 0.001; Figure 2).

### Gene family evolution

*Orthofinder2* grouped protein sequences from 18 Myodontine rodents into a total of 23,020 orthogroups. After removing orthogroups with either high or no variation in the number of genes across species, the dataset was reduced to 21,347 orthogroups. On average, all species included in our analysis showed gene family contraction, albeit of varying magnitude. *Mus musculus* had the highest number of significant changes (p < 0.01) in gene family size (92 orthogroups), while *Dipodomys ordii* had no significant changes (Figure 3). The number of gene families with significant contractions or expansions varied between 16 and 24 among *Peromyscus* species, with more expansions than contractions except in the cactus mouse, which exhibited four gene family expansions and 12 contractions (Figure 3). Four of these gene families contained genes associated with sperm motility, four included ribosomal proteins, three were associated with immune response, and two included genes in the ubiquitin-like (ubl) conjugation pathway. Other functions included pheromone reception, cytoskeletal protein binding, and prohibitin (one gene family each).

**Figure 3.**
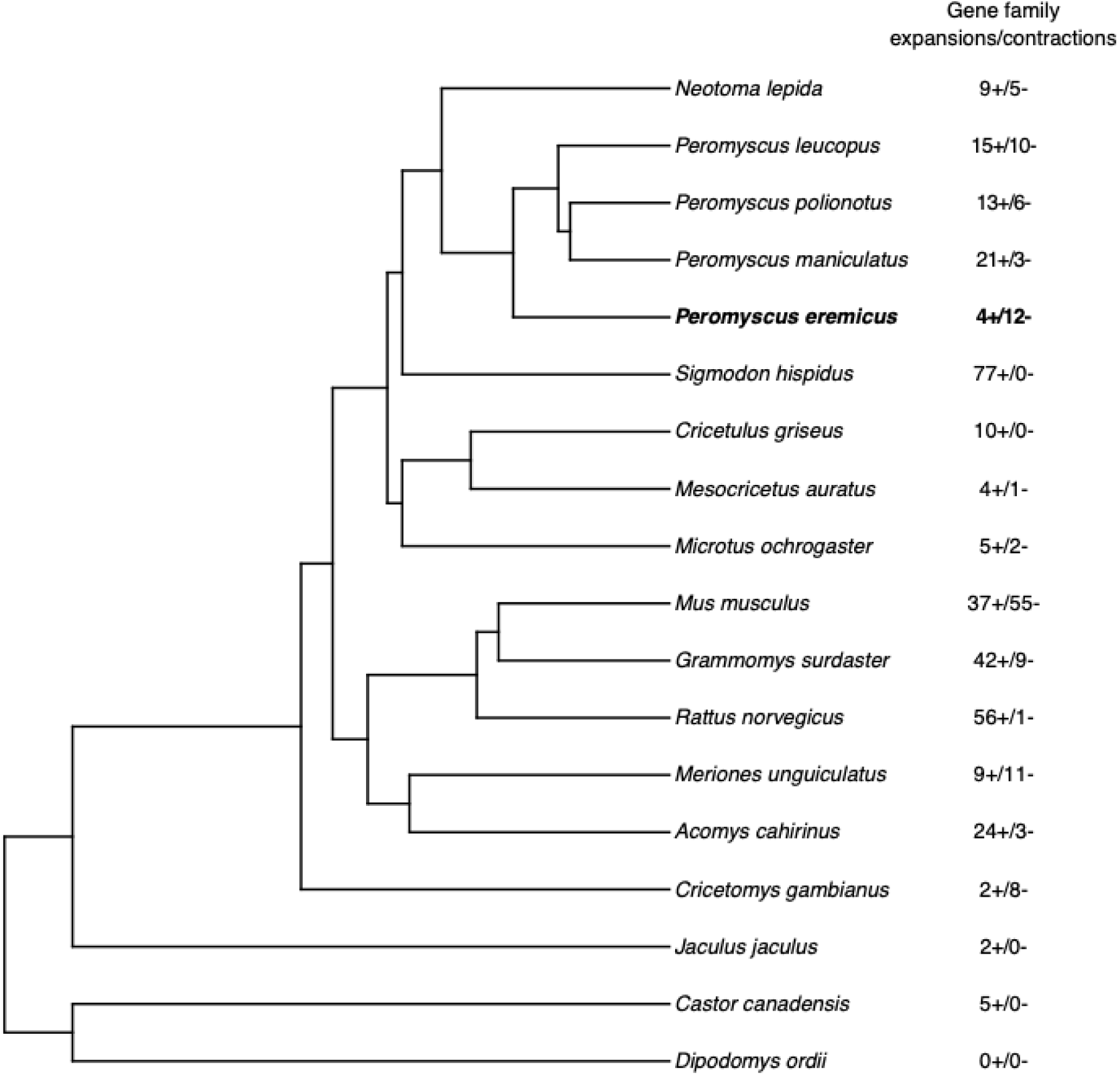
Species tree built in *Orthofinder2* from protein sequences of 18 species in the Myodonta clade (Order: Rodentia). Beside each species name are the number of gene families that underwent significant (p < 0.05) expansions (+) or contractions (-).

### Identification of selective sweeps

Analysis of the signatures of selective sweeps yielded a total of 232 sites under selection. Of these, 119 clustered in 44 larger regions that included two or more adjacent CLR outliers (Supplementary Figure 4). By retrieving the genes closest to each peak (one in both up- and down-stream directions), we compiled a list of 186 genes associated with selective sweeps (Supplementary Table 3). Fourteen of these genes, including many putative olfactory and vomeronasal receptors, were not matched with a corresponding GO term. Ribosomes were overrepresented among ‘cellular components’, with eight GO terms pointing to this organelle (p < 0.001, after Bonferroni correction). In addition to this, 279 biological processes and 89 molecular functions were significantly overrepresented (before correction for multiple tests; full list in Supplementary Table 4 and 5, respectively). GO terms clustering in *REVIGO* showed that terms with the lowest p-values under ‘biological processes’ included ‘membrane organization’, ‘cellular amide metabolism process’, ‘translation’, ‘ribosome assembly’, and ‘detection of chemical stimulus involved in sensory perception of bitter taste’ (Figure 4, Supplementary Table 3); while under ‘molecular functions’ they included ‘structural constituent of ribosome’, ‘binding’, ‘olfactory receptor binding’, and ‘mRNA binding’ (Figure 4, Supplementary Table 4).

**Figure 4.**
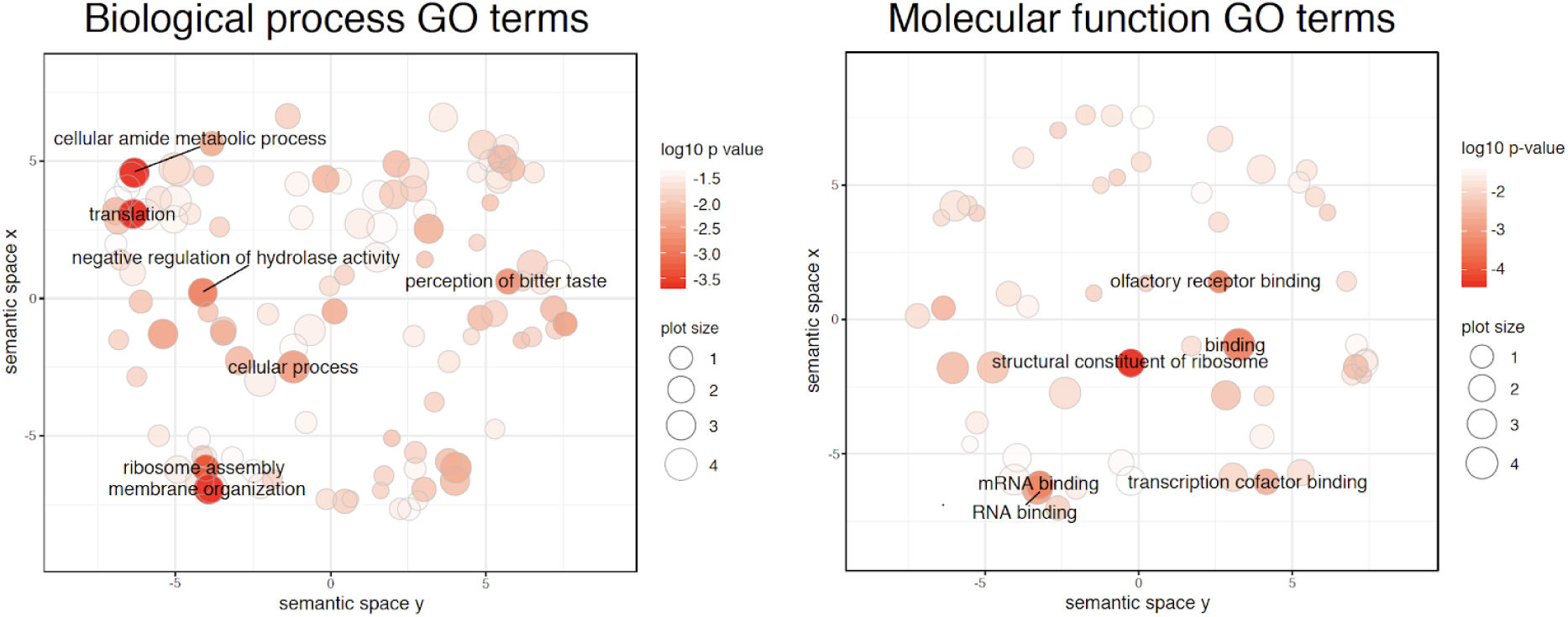
Scatterplot showing clusters representative of enriched GO terms after semantic reduction in *REVIGO* for biological process GO terms (left) and molecular function GO terms (right). Only the names of GO clusters with a p-value < 10^-2.5^ are shown for visual clarity. The full list of genes and reduced GO terms in *REVIGO* can be found in Supplementary Tables 3, 4 and 5).

Contrary to predictions, mean π and Tajima’s D were significantly higher across the candidate areas for selective sweeps when compared to genome-wide means (Wilcoxon test, p < 0.001 for both π and Tajima’s D; Figure 5). Among the candidate genes from previous studies, 8 (all *Cyp4* genes, *SLC8A1*, and *aqp4*, *aqp5*, *aqp8,* and *aqp12*) and 6 genes (the *Cyp4a* gene cluster, *Cyp4v2*, *SLC8A1*, and *aqp5*, *aqp9,* and *aqp12*) showed significant deviations from genome-wide average in π and Tajima’s D, respectively (p < 0.05 after Benjamini-Yekutieli correction for multiple testing), but not always in the predicted direction (Supplementary Figure 5). Among the *Cyp4* genes, *Cyp4f* showed a modest decrease in π; among aquaporins, *aqp8* showed a decrease in π, and *aqp9* showed a decrease in Tajima’s D; and *SLC8a1* showed the greatest reduction in π and Tajima’s D overall (Supplementary Figure 5).

**Figure 5.**
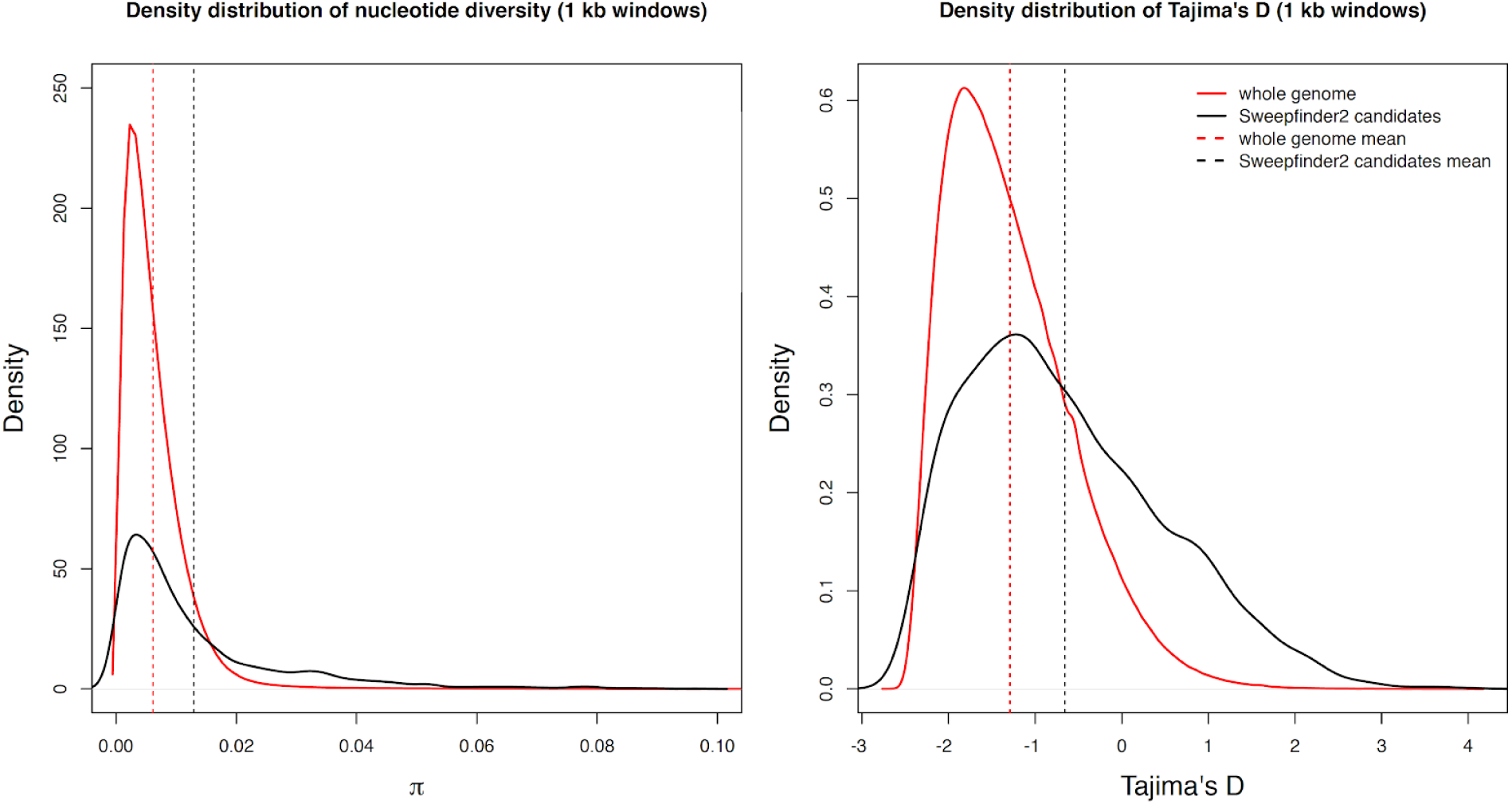
Density plots comparing distribution of π (left) and Tajima’s D (right) across the genome (in red) and across *Sweepfinder2* candidate regions only (in black). Values are calculated in 1 kb non-overlapping windows along the genome. Dashed vertical lines show the means across the genome in red and across *Sweepfinder2* candidate regions only in black. Means across the genome and across *Sweepfinder2* candidate regions only are highly significant in both cases (p < 0.001).

Number of sweeps in each chromosome did not correlate with mean π (p > 0.05). A correlation with either chromosome size or number of indels (p < 0.01) was entirely driven by the outlier behaviour of chromosome 1, and it did not hold when chromosome 1 was removed from the dataset (p > 0.05).

## Discussion

### A chromosome-level assembly for the cactus mouse

A high-quality chromosome-level assembly of the cactus mouse genome allowed us to investigate genomic patterns of variation, differentiation, and other genomic features (i.e. genes, repetitive elements, number and size of chromosomes), and to identify regions of the genome that may be associated with desert adaptations. As the number of publicly available *Peromyscus* genome assemblies increases (Colella et al., 2019), the cactus mouse genome will provide additional insights into adaptation, speciation and genome evolution when analyzed in a comparative framework. For example, our comparison of the cactus mouse and the deer mouse genomes revealed higher than expected genome stability considering the divergence time between the two species (∼9 MYA), and their large effective population sizes and short generation times (Bromham, 2009; Charlesworth, 2009). Our synteny analysis confirms at the sequence level what is reported from karyotype analyses of several *Peromyscus* spp. (Smalec et al., 2019), i.e. that between *P. eremicus* and *P. maniculatus* there is no variation in chromosome number and no interchromosomal rearrangements but abundant intrachromosomal variation. The genome stability among *Peromyscus* species (Long et al., 2019) is in sharp contrast to the Muridae family (Order Rodentia) among *Mus* species and the brown rat (*Rattus norvegicus*), where the number of chromosomes varies dramatically, even within the *Mus* genus, and large chromosomal rearrangements are abundant (Thybert et al., 2018).

### High genetic diversity and differentiation

Our results indicate that population differentiation between cactus mouse populations inhabiting the Motte and Deep Canyon Reserves in Southern California is high despite being separated by only 90 km. Patterns of differentiation across the genome were high without distinguishable F_ST_ peaks (Figure 1B). The distribution of allele frequency changes is suggestive of prolonged geographical isolation, which is consistent with the Peninsular Ranges mountains acting as a dispersal barrier. Although this is not surprising given the limited dispersal ability of cactus mice, the close proximity of these two sampling locations suggests that population structure across the species range, which spans more than 2500 km from Nevada (USA) to San Luis Potosì (Mexico), is likely to be strong. Previous analyses based on a single mitochondrial marker split the *P. eremicus* species complex into three species – *P. eva*, *P. fraterculus* and *P. merriami* – plus West and East *P. eremicus* clades (Riddle et al., 2000). Our whole genome analyses suggest that population structure could be pronounced even within each *P. eremicus* clade, which warrants further investigation to elucidate the taxonomic status of these species and to reveal potential differences in adaptation to local desert conditions.

As standing genetic variation is the main source of adaptive genetic variation (Barrett and Schluter, 2008), characterizing levels and distribution of sequence and structural variation can help understand how and where in the genome adaptations evolve. With more than 43 million high-quality SNPs and ∼87,000 indels, the cactus mouse exhibits high levels of standing genetic variation, which is consistent with large effective population sizes and comparable to what has been reported for the congeneric white-footed mouse (*P. leucopus*; 42 million SNPs across 26 wild-caught individuals; (Long et al., 2019). While SNPs are the main, and most often the only, type of variation screened in genomic studies of adaptation (Wellenreuther et al., 2019), here we show that small- to mid-sized indels, a type of structural variant, are common and cover ∼4% of the genome. Given that our analysis of structural variation leveraged sequence data from only one individual, the level of standing structural variation in the population is likely much higher. Pezer et al. (2015) found that indels covered ∼2% of an individual wild house mouse (*M. musculus domesticus*) genome compared to a reference genome, likely an underestimation considering that the analysis was based on variation of read depth only. The full characterization of standing genetic variation in a wild fish showed that levels of structural variation were threefold the levels of sequence variation (SNPs) across 12 individuals sequenced at high coverage (Catanach et al., 2019). To this end, while the inclusion of our high coverage 10X Genomics dataset allowed us to characterize structural variation in a single individual, future work conducted at the population level will allow us to integrate more deeply the role of structural variation in the evolution of desert adaptation in the cactus mouse.

Nucleotide variation and number of indels were significantly correlated with chromosome size in the cactus mouse. With increasing chromosome length, sequence variation decreased while the number of indels increased. While a positive, linear relationship between chromosome size and number of indels is expected, a negative correlation between chromosome size and nucleotide diversity may be explained by recombination rate, which is higher in shorter chromosomes (Kaback et al., 1992). Chromosome 1 was an outlier compared to the rest of the autosomes in that it was the largest (203 Mb), with the greatest difference from the next largest autosome (26.6 Mb, range 26.6 Mb - 561 Kb), and it had higher nucleotide diversity than expected based on size (i.e. similar to what is expected for much smaller chromosomes). The outlier behavior of chromosome 1 could be due to an ancient fusion of two smaller chromosomes demonstrated by syntenic analysis of *Peromyscus leucopus* and *Mus musculus* (Long et al., 2019).

Sex chromosomes generally harbor lower nucleotide diversity (Wilson Sayres, 2018) and higher differentiation (Presgraves, 2018) when compared to autosomes due to their reduced effective population size, different mode of inheritance, and their role in the evolution of reproductive barriers. We did observe lower π in comparison to autosomes (45-71% of mean π for each autosome), slightly lower than neutral expectations assuming equal sex ratio (75%). Demographic processes and/or selection could be implicated in this additional reduction, but their relative roles were not tested here. However, contrary to our expectations, analysis of genomic differentiation based on F_ST_ showed that the X chromosome was less differentiated than the autosomes in the interpopulation comparison. In fact, sex chromosomes are more differentiated than autosomes in 95% of the studies for which this information is available (Presgraves, 2018). In two cases regarding mammals, domestic pigs and wild cats, lower differentiation on the X chromosomes was ascribed to hybridization and introgression (Ai et al., 2015; Li et al., 2016). Hybridization has been reported among several *Peromyscus* species (Barko and Feldhamer, 2002; Leo and Millien, 2017), and it represents a viable hypothesis given that the cactus mouse is sympatric with the canyon mouse (*Peromyscus crinitus*) and numerous other *Peromyscus* species throughout much of its range. Population genomic data from additional *Peromyscus* species are necessary to assess how common this pattern is within and across species, to test the hybridization hypothesis, to identify potential donor and recipient species, and to test the potential role of hybridization and introgression in desert adaptation.

Neither sequence nor structural variation at the chromosome level was a strong predictor of the number of selective sweeps in a chromosome. However, mean π and Tajima’s D were significantly higher in the areas affected by selective sweeps than across the whole genome. Theory predicts that a selective sweep should remove variation from the adaptive site and its surroundings, thus resulting in a localized reduction in π and lower Tajima’s D relative to the ancestral level of variation (Kim and Stephan, 2002; Smith and Haigh, 1974). However, the signature of a selective sweep, and the ability to detect it, depends on the strength of selection, the recombination rate around the selected site, and whether the sweep is hard, soft, or incomplete (i.e. whether a single or multiple haplotype carries the beneficial allele, or the allele hasn’t reached fixation yet; Messer and Neher, 2012). *Sweepfinder2* is best suited to detect hard selective sweeps and has limited power to identify soft or incomplete sweeps (DeGiorgio et al., 2016; Huber et al., 2016; Nielsen et al., 2005). Therefore, if we do not observe a general reduction of diversity compared to the genome-wide average around adaptive sites, it could be for one or a combination of the following reasons. a) High recombination rates due to a large effective population size may break up linkage among neighboring sites, thus reducing the size of the typical, diagnostic dip in diversity to a point of non-detectability. b) Due to computational limitations (*Sweepfinder2* is not able to parallelize, leading to long run times) we estimated the CLR of a site every 10 kb, which may not be dense enough to pinpoint the exact location of the sweep and reveal narrow reductions in π. c) If soft sweeps are indistinguishable from hard sweeps when selection is strong (Harris et al., 2018), selective sweeps may preferentially occur in areas of high standing genetic variation. Finally, d) even if a reduction in diversity occurs relative to ancestral levels, π may still not drop under the genome-wide average, especially if diversity was originally high. On chromosome 9, for example, where three consecutive outlier regions extend over 440 kb, the reduction of π and Tajima’s D was drastic, suggesting that coarse resolution may prevent the detection of diversity dips around selected sites if the genomic area affected is small.

### Lost in translation: pervasive signature of selection in genes associated with protein synthesis and degradation

Together, the analyses of gene family evolution and selective sweeps strongly indicate that traits associated with the synthesis and degradation of proteins have evolved under the influence of natural selection. Four ribosomal protein families are either expanded or contracted, and gene ontology analysis demonstrated an enrichment of terms associated with ribosomes (e.g., ribosome assembly and translation, structural constituents of ribosomes, mRNA binding, and unfolded protein binding). We also report a significant contraction of a gene family associated with the ubl conjugation pathway, which was similarly identified in an analysis of selective sweeps. Ubiquitin and ubl-proteins function either as a tag on damaged proteins to be lysed or as regulators of interactions among proteins (Hochstrasser, 2009). Cactus mice face many stressors including high temperatures and lack of water. Heat causes cellular stress directly, via thermal stress, and indirectly, by exacerbating the negative effects of dehydration due to lack of water and rapid water loss (e.g., respiratory water loss, evaporative cooling). Thermal and hyperosmotic stress can suppress the transcription and translation machinery, increase DNA breaks and protein oxidation, and cause cell cycle arrest, and eventually apoptosis and cell death (Burg et al., 2007; Kampinga, 1993). However, the strongest and most immediate effect of thermal and hyperosmotic stress is protein denaturation (Burg et al., 2007; Kampinga, 1993; Lamitina et al., 2006). Our results are consistent with the expected cellular response to both thermal and hyperosmotic stress, which have similar physiological effects even though the underlying mechanisms may differ. A meta-analysis of genomics and transcriptomics studies investigating the evolutionary response to different thermal environments in metazoans, including invertebrates to mammals, highlighted ‘translation’, ‘structural constituents of ribosomes’, and ‘ribosome’ as the gene ontology terms most commonly enriched (Porcelli et al., 2015), in line with our results. Similarly, many genes mediating the cellular response to hyperosmotic stress are involved in the regulation of protein translation and the elimination of denatured proteins in *Caenorhabditis elegans* (Lamitina et al., 2006). These analyses suggest that selection has acted strongly on genes responsible for protection against thermal and/or hyperosmotic stress or for efficiently removing damaged proteins and resuming translation after acute stress (Kampinga, 1993). Additionally, as the volume of dehydrated cells decreases causing rearrangements in the cytoskeleton (Burg et al., 2007), the significant contraction of a cytoskeletal protein gene family could also point to additional adaptations to hyperosmotic stress in the cactus mouse. Acute dehydration experiments on captive cactus mice also found limited tissue damage and apoptosis in the kidneys of dehydrated individuals (MacManes, 2017), consistent with our hypothesis. Negative regulation of cell death was one of the most significant GO terms, suggesting that these genes may be under selection to avoid tissue necrosis during acute or chronic stress.

### Life in the desert involves dietary and metabolic adaptations

The GO analysis of genes associated with selective sweeps indicated an enrichment for bitter taste receptors. The perception of bitter taste has evolved to allow organisms to avoid toxic compounds found in many plants and insects (Garcia and Hankins, 1975; Glendinning, 1994). Although herbivorous and insectivorous animals generally have a larger repertoire of bitter taste receptors compared to their carnivorous counterparts (Li and Zhang, 2014; Wang and Zhao, 2015), they are also less sensitive to bitterness (Glendinning, 1994). The cactus mouse is omnivorous, with a diet predominantly based on seeds, insects, and green vegetation with proportions varying according to seasonal availability (Bradley and Mauer, 1973; Meserve, 1976). We hypothesize that repeated signal of selective sweeps at bitter taste receptor genes may have increased the frequency of alleles that decrease bitter sensitivity, thus making a greater variety of food palatable to the cactus mouse in an environment that is characterized by scarcity of resources and an abundance of bitter-tasting plants and insects.

Chromosome 9 showed the largest and strongest selective sweep in the genome (Supplementary Figure 4). This area was associated with *Gdf10* (growth/differentiation factor 10), the only annotated gene of known function in the region, which is involved in osteogenesis and adipogenesis. Overexpression of *Gdf10* in the adipose tissues of mice prevents weight gain under a high-fat diet and affects their metabolic homeostasis, including oxygen consumption and energy expenditure (Hino et al., 2017). The drastic loss of weight and the starvation-like response reported in experimentally dehydrated cactus mice suggests that lipid metabolism has a role in the adaptive response to dehydration (MacManes, 2017). Experimental water deprivation induced higher food consumption and loss of body fat in the spinifex hopping mouse (*Notomys alexis*), a desert-specialist rodent (Takei et al., 2012). In camels, accelerated evolution of genes associated with lipid metabolism was associated with food scarcity in the desert (Wu et al., 2014). The strong signature of selection around *Gdf10* in the cactus mouse therefore warrants further investigation, as adaptive changes in lipid metabolism may be pivotal for survival in the desert.

Among the candidate genes we selected from previous studies, only *SLC8a1* – the sodium carrier gene – show significant reduction in both π and Tajima’s D, consistent with a selective sweep. However, *Cyp4v2* – one of the genes in the arachidonic acid pathway – was in proximity of a selective sweep on chromosome 17. This gene shows similar catalytic properties to other *Cyp4* genes in the arachidonic acid pathway and is commonly expressed in retinal, kidney, lung, and liver of humans (Nakano et al., 2009). Known to be strongly associated with ocular disease, its role, however, has not been investigated in the context of water-sodium balance in the kidneys. The discrepancy between the strong changes in gene expression in *Cyp4* genes between hydrated and dehydrated mice (MacManes, 2017) and the results presented here suggest that these genes affect kidney physiology predominantly via gene expression and potentially through changes in regulatory regions that we have not targeted explicitly. Future comparative analyses of sequences from additional rodents, including other desert-adapted species, will help us understand the relative role of gene expression regulation versus coding changes in adaptation to desert environments.

Comparisons with studies on other desert-adapted mammals highlight a combination of convergent and idiosyncratic adaptations to life in the desert. Although the arachidonic acid pathway showed signatures of selection in camels, sheep, and the cactus mouse (Bactrian Camels Genome Sequencing and Analysis Consortium et al., 2012; MacManes, 2017; Yang et al., 2016), the specific genes involved differed among species and between the mechanisms by which they putatively affected adaptive phenotypes: gene family contractions and expansions in camel, selective sweeps in desert sheep, and changes in gene expression under acute dehydration in the cactus mouse. In addition to the stress imposed by heat and lack of water, comparative genomics analyses suggest that camels have unique adaptations to avoid the deleterious effects of dust ingestion and intense solar radiation (Wu et al., 2014). The cactus mouse avoids solar radiation altogether with a nocturnal lifestyle and shows strong signatures of selection at receptors for bitter taste, consistent with adaptation to a diet based on bitter-tasting desert plants and insects. Desert woodrats (*Neotoma lepida*) are highly specialized to bitter and toxic plants, such as juniper (*Juniperus monosperma*) and creosote bush (*Larrea tridentata*), and have evolved several adaptations to consume them, including detoxifying gut microbiomes (Kohl et al., 2014) and hepatic enzymes (Skopec and Dearing, 2011). Although the repertoire of bitter taste receptors, or their sensitivities to bitter taste, has not been investigated in desert rodents, nor our genomic analyses of cactus mice highlighted detoxification genes, it is evident that many desert species have evolved different strategies to cope with bitter, toxic plants in the absence of more palatable options.

### Signatures of selection potentially involved in reproductive isolation

We reported significant evolutionary changes linked to genes for sperm motility, spermatogenesis, and pheromone reception that may lend support to a role of sexual selection in the evolution of the cactus mouse genome. The comparison of sperm morphology and behaviour in *Peromyscus* species has revealed a link between sperm traits and reproductive strategies. Sperm of the promiscuous *P. maniculatus*, for example, can aggregate on the basis of relatedness, thus increasing motility and providing a competitive advantage against other males, whereas sperm of the monogamous *P. polionotus* lacks these adaptations (Fisher et al., 2014; Fisher and Hoekstra, 2010). The four gene families associated with sperm motility showed a conspicuous contraction in the cactus mouse when compared to other *Peromyscus* species (9 versus 26, 17 and 21 in *P. leucopus*, *P. maniculatus*, and *P. polionotus*, respectively). Although these results may suggest a correlation between reduction in the number of sperm motility genes and a monogamous reproductive strategy, *P. maniculatus* also has fewer genes than *P. polionotus* in these gene families (17 versus 21 in total). Nonetheless, these candidate genes represent interesting targets for future studies on sperm competition within the cactus mouse and among *Peromyscus*.

## Conclusions

The high-quality assembly of the cactus mouse genome and the candidate genes identified in this study build on the growing body of genomic resources available to further understand the genomic and physiological basis of desert adaptation in the cactus mouse and other species. Taken together, our results indicate that the strongest signatures of selection in the cactus mouse genome are consistent with adaptations to life in the desert, which are mostly, but not solely, associated with high temperatures and dehydration. Contrary to expectations, we did not find a pervasive signature of selection at genes involved in water-sodium balance in the kidneys. However, this does not necessarily minimize the relative role of these organs under thermal and hyperosmotic stress, as we have not yet tested when and where in the body the expression of these candidate genes is beneficial. Our analyses also show that signatures of selection are widespread across the cactus mouse genome, with all autosomes showing selective sweeps, and that they are not affected by chromosome-level patterns of standing genetic variation, sequence or structural. Sweeps seem to be associated with high local π, instead. Dynamic gene families and enrichment of several GO terms associated with selective sweeps indicate that the genetic basis of at least some desert-adapted traits may be highly polygenic. In the future, evolutionary and physiological genomics work stemming from these results will allow us to better characterize the phenotypes and genotypes associated with desert adaptations in the cactus mouse, and to understand how they evolved.

## Methods

### Ethics Statement

All sample collection procedures were approved by the Animal Care and Use Committee located at the University of California, Berkeley (2009 samples, protocol number R224) and University of New Hampshire (2018 samples, protocol number 130902) as well as the California Department of Fish and Wildlife (permit number SC-008135) and followed guidelines established by the American Society of Mammalogy for the use of wild animals in research (Sikes and Animal Care and Use Committee of the American Society of Mammalogists, 2016).

### Genome assembly and annotation

We extracted DNA from the liver of a female cactus mouse (ENA sample ID: SAMEA5799953) captured near Palm Desert, CA, USA using a Qiagen Genomic Tip kit (Qiagen, Hilden, Germany). We built two short-insert Illumina libraries (300 bp and 500 bp inserts) using an Illumina Genomic DNA TruSeq kit, following manufacturer recommendations. For scaffolding, we added four mate pair libraries (3 kb, 5 kb, 7 kb, 8 kb) prepared using a Nextera Mate Pair Library Prep kit (Illumina, San Diego, CA, USA). Each library was sequenced on an Illumina HiSeq 2500 sequencer by Novogene (Sacramento, CA, USA) at a depth of approximately 30x for each short-insert library, and 5x for each mate pair library. After adapter trimming, libraries were assembled using the program *ALLPATHS* (Butler et al., 2008). The resulting assembly was gap-filled using a PacBio library (Pacific Biosciences of California, Inc., Menlo Park, CA, USA), constructed from the same DNA extraction and sequenced at ∼5x coverage, using *PBJelly* (English et al., 2012). We error-corrected the resulting assembly with short-insert Illumina data and the *Pilon* software package (Walker et al., 2014).

To improve the draft assembly, we used proximity-ligation data (Hi-C) to further order and orient draft scaffolds. We prepared a Hi-C library using the Proximo Hi-C kit from Phase Genomics (Seattle, WA, USA). We used ∼200 μg of liver from a second wild-caught animal and proceeded with library preparation following the protocol for animal tissues. The Hi-C library was sequenced at Novogene using one lane of 150 bp paired-end reads on an Illumina HiSeq 4000 platform. To arrange the draft scaffolds in chromosomes we used the program *Juicer* in an iterative fashion (Durand et al., 2016b). Following each run, we loaded the *.map* and *.assembly* files generated by *Juicer* into *Juicebox* (Durand et al., 2016a), the accompanying software developed to visualize crosslinks, and corrected misassemblies manually. We ran *Juicer* until no well-supported improvements in the assembly were observed. We thus obtained 24 chromosome-sized scaffolds plus 6,785 unplaced short scaffolds. We calculated assembly statistics with the *assemblathon_stats.pl* script from the Korf Lab (https://github.com/KorfLab/Assemblathon/blob/master/assemblathon_stats.pl) and assessed assembly completeness with *BUSCO v3* (Simão et al., 2015) and the Mammal gene set.

To standardize chromosome naming and enable future comparative analyses, we used the genome assembly of the deer mouse (*Peromyscus maniculatus bairdii*; NCBI Bioproject PRJNA494228) to name and orient the cactus mouse chromosomes. We used *mummer4* (Marçais et al., 2018) to align the cactus mouse genome to the deer mouse genome with the function *nucmer* and the options *–-maxgap 2000* and *–minclust 1000*. We filtered alignments smaller than 10 kb with *delta-filter* and plotted the alignment using *mummerplot*. These genome alignments also allowed us to test for synteny between the two species, diverged ∼9 million years ago (timetree.com), and to assess the degree of structural divergence between the two genomes.

We identified transposable elements and other repetitive elements using *RepeatMasker v.4.0* (Smit et al., 2015) and the Rodentia dataset. The masked genome was annotated using *Maker v.2.3.1* (Cantarel et al., 2008) and the *Mus musculus* reference protein dataset.

### Whole genome resequencing

We sequenced the genomes of an additional 26 cactus mice collected from two locations in Southern California: Motte Rimrock and Boyd Deep Canyon Reserves (both belonging to the University of California Natural Reserve System; Table 1). The Motte Rimrock Reserve (Motte hereafter) sits on a broad, rocky plateau and supports both coastal and desert habitats as it is located equidistant between the Pacific coast and the Colorado Desert. We captured cactus mice in xeric areas characterized by rocky outcrops, which constitutes their typical habitat. The Boyd Deep Canyon Reserve (Deep Canyon hereafter) is a large natural reserve extending from low to high elevation (290-2657 m). We sampled at two locations in the lower elevation part of the Deep Canyon Reserve: Deep Canyon (290 m a.s.l.), the driest and hottest location, with average monthly temperatures between 10-40 °C and mean annual rainfall of 15 cm, and Agave Hill (820 m a.s.l.), with average monthly temperatures between 9-35 °C and mean annual rainfall of 18 cm. Samples from Deep Canyon were collected in 2009 and 2018 (Table 1).

Genomic libraries were prepared at the Biotechnology Resource Center at Cornell University (Ithaca, NY, USA) using the Illumina Nextera Library Preparation kit and a modified protocol for low-coverage whole genome resequencing (‘skim-seq’). Individually barcoded libraries were sequenced at Novogene using 150 bp paired-end reads from one lane on the Illumina NovaSeq S4 platform. We conducted an analysis of sequencing read quality and trimmed adapters from raw sequencing data with *fastp* (Chen et al., 2018). We mapped sequences from each of the 26 individuals the cactus mouse reference genome using *bwa mem* (Li and Durbin, 2009) and removed duplicates with *Samblaster* (Faust and Hall, 2014). The resulting BAM files were sorted and indexed using *Samtools* (Li et al., 2009).

As sequencing depth was variable among individuals (raw coverage between ∼2-17X), we called variants in *ANGSD* (Korneliussen et al., 2014) as its algorithm takes into account genotype uncertainty associated with low-coverage data. To identify a list of high-confidence variable sites, we ran a global variant calling analysis including all 26 individuals. To be included in our analyses, a site had to satisfy the following criteria: p-value below 10^-6^, minimum sequencing and mapping qualities above 20, minimum depth and number of individuals equal to half of the number of individuals included in the analysis (13 out of 26), and a minor allele frequency (MAF) above 1%.

### Genomic differentiation across space and time

To estimate the effects of temporal and spatial distance on levels of genomic differentiation among individuals, we first ran an Analysis of Principal Components (PCA) of genetic data using the *ngsCovar* program from *ngsTools* (Fumagalli et al., 2014). We used all high-quality SNPs (defined above) called at the species level and ran the PCA using genotype posterior probabilities, rather than called genotypes, as input. To estimate genetic distance between individuals controlling for the effect of varying depth of sequencing across individuals, we downsampled to a single base for each site included in the high-quality SNPs list and performed multidimensional scaling analysis (MDS) in *ANGSD*. Preliminary analysis revealed the presence of an individual from Motte that grouped with neither population. This individual was subsequently removed and variants were re-called.

After outlier exclusion, we reran *ANGSD* to estimate allele frequencies in Motte and Deep Canyon separately. We provided a list of high-confidence SNPs for use in downstream analyses and applied the same filters as in the global SNP calling, excluding SNP p-value and MAF thresholds, with the major allele fixed across runs. We used the sample allele frequencies (.mafs file) from each population to calculate the 2D Site Frequency Spectrum (SFS), which we also used as prior for estimating F_ST_, a measure of genetic differentiation. We calculated average F_ST_ across autosomes and across the X chromosome separately, and investigated patterns of differentiation across the genome in 50 kb sliding windows.

### Sequence and structural standing genetic variation

To estimate levels and patterns of standing genetic variation within the cactus mouse, we analyzed samples from both populations together. We calculated the overall proportion of polymorphic sites and nucleotide diversity (π) in 50 kb sliding windows. Estimates of π for each polymorphic site were based on the maximum likelihood of the SFS calculated with *realSFS* in *ANGSD*. To obtain accurate estimates of diversity, we corrected global and window estimates of π by the number of variant and invariant sites covered by data. We estimated genome coverage, total and per window, by rerunning *ANGSD* (including all 25 samples) using the same filtering parameters we used for the global calling variant but without the SNP p-value and the MAF filters. We then divided the sum of per-site π by the number of variant and invariant sites in a given window.

To investigate the distribution of structural variation in the cactus mouse genome, we sequenced the genome of an additional individual from the Deep Canyon site within the Deep Canyon Reserve (sampled in 2009) using 10X Genomics (Pleasanton, CA, USA). This method is based on linked-reads technology and enables phasing and characterization of structural variation using synthetic long-range information. We used the program *Long Ranger v.2.2.2* from 10X Genomics and ran it in whole genome mode with *longranger wgs* using *Freebayes* (Garrison and Marth, 2012) as variant caller. Finally, we tested whether chromosome size was a predictor of either sequence or structural standing variation using linear models in *R* (R Core Team 2019).

### Gene family evolution in the cactus mouse

To investigate gene family contractions and expansions in the cactus mouse, we analyzed the genomes of 25 additional species (Supplementary Table 1) within the Myodonta clade (Order: Rodentia), which includes rats, mice, and jerboas, for which genome assemblies were publicly available from NCBI in June 2019. To avoid potential biases due to different gene annotation strategies, we re-annotated all 25 genomes with the same strategy used for the cactus mouse (see above). Genome quality was evaluated using *BUSCO*. We identified groups of orthologous sequences (orthogroups) in all species using the package *Orthofinder2* (Emms and Kelly, 2018) with *Diamond* as protein aligner (Buchfink et al., 2015). In a preliminary run, we observed that fewer orthogroups and fewer genes per orthogroup were identified in species with lower genome assembly quality. Therefore, we filtered assemblies that had less than 70% complete benchmarking universal single-copy orthologs (BUSCOs) and thus retained 18 species (Supplementary Table 1).

We analyzed changes in gene family size accounting for phylogenetic history with the program *CAFE v4.2.1* (De Bie et al., 2006). We filtered both invariant orthogroups and those that varied across species by more than 25 genes. We used the rooted species tree inferred by *Orthofinder2* and ran the analysis using a single value for the death-birth parameter (λ) estimated in *CAFE* for the whole tree. Finally, we summarized the results with the python script *cafetutorial_report_analysis.py* from the *CAFE* developers (https://hahnlab.github.io/CAFE/manual.html).

### Detection of signatures of selection

We used an integrative approach to detect signatures of selection from the population genomics data. First, we used *Sweepfinder2* (DeGiorgio et al., 2016; Nielsen et al., 2005) to detect recent selective sweeps. We ran these analyses using all 25 individuals, based on the rationale that potential differences between the two populations (Deep Canyon and Motte) due to local adaptation or demography would be swamped by signatures common in the species across populations. As recommended by Huber et al. (2016), we included both variant and invariant sites, but we could not reconstruct the ancestral state of these sites due to lack of data from closely related species. The X chromosome was excluded from this analysis. We converted allele frequencies estimated in *ANGSD* to allele counts, and estimated the SFS from the autosomes only in *SweepFinder2*. We then tested for sweeps using *SweepFinder2* with the *-l* setting, i.e. using the pre-computed SFS, and calculated the composite likelihood ratio (CLR) and α every 10,000 sites. Only the peaks with CLR values above the 99.9^th^ percentile of the empirical distribution of CLR values were considered under selection. We functionally annotated the closest genes to these outlier peaks and ran a Gene Ontology (GO) enrichment analyses to test whether genes under putative selection were enriched for a particular function or pathway. We performed this analysis on the geneonotology.org webpage using the program *Panther* (Mi et al., 2017) and the *Mus musculus* gene set as reference. As GO terms are hierarchical, we summarized these results with the software package *REVIGO* (Supek et al., 2011), which uses a clustering algorithm based on semantic similarities, setting the similarity threshold at 0.5.

Finally, we generated a list of *a priori* candidate genes potentially involved in desert adaptation. These included the genes that were most differentially expressed in response to experimental dehydration, including 11 *Cyp4* genes from the arachidonic acid metabolism pathway, and the sodium carrier gene *Slc8a1* (MacManes, 2017). We also included the 9 aquaporins, which are important in water reabsorption in the kidney but were not differentially expressed in hydrated versus dehydrated cactus mice (MacManes, 2017). We integrated this list of candidate genes with the genomic areas showing strong signatures of selective sweeps in the *Sweepfinder2* analysis. As decreases in π and Tajima’s D can also be indicative of selective sweeps, we calculated these two statistics in 1 kb windows across these candidate regions, plus an additional 10 kb flanking on each side.

## Data availability

All read data for the genome assembly are housed on ENA under project ID PRJEB33593. Specifically, genome assembly (ERZ1195825), 300 bp PE (ERR3445708), 500 bp PE (ERR3446161), 8 kb mate pair (ERR3446162), 5 kb mate pair (ERR3446317), 3 kb mate pair (ERR3446318), 7 kb mate pair (ERR3446319), Hi-C (ERR3446437), 10X Genomics (ERR3447855). Whole genome resequencing data for 26 cactus mice are housed on ENA under project ID PRJEB35488. Scripts for the genome assembly and all other analyses can be found at https://github.com/atigano/Peromyscus_eremicus_genome/

## Acknowledgements

We would like to thank Christopher Tracy for access to the Boyd Deep Canyon Reserve, Adam Stuckert and Douglas Kelt for help with field sampling, the Biotechnology Resource Center at Cornell University for preparation of the whole genome resequencing libraries, the MacManes Lab and the Evolutionary Genomics Journal Group at the University of New Hampshire (UNH) for useful comments on earlier versions of the manuscript. All analyses were performed on the UNH Premise Cluster. This work was funded by the National Institute of Health National Institute of General Medical Sciences to MDM (1R35GM128843)

## Author contributions

AT and MDM conceived the study, collected the data, assembled the cactus mouse genome and performed analyses. JPC provided input on the interpretation of results. AT wrote the first version of the paper and AT, JPC and MDM reviewed and edited the paper.

