## Supplementary Figures and Tables for "Comparative and population genomics approaches reveal the basis of adaptation to deserts in a small rodent"

**Supplementary Table 1.** List of species in the Myodonta clade (Order Rodentia) included in the gene family evolution analysis and associated genome completeness statistics measured in proportion of complete and missing BUSCOs.

| Species | Complete BUSCO % | Missing BUSCO % | Retained (> 70% complete BUSCOs) |
|---|---|---|---|
| *Acomys cahirinus* | 75.70 | 7.30 | X |
| *Castor canadensis* | 87.00 | 5.70 | X |
| *Cricetomys gambianus* | 87.00 | 3.50 | X |
| *Cricetulus griseus* | 93.00 | 3.10 | X |
| *Dipodomys ordii* | 94.80 | 2.20 | X |
| *Grammomys surdaster* | 94.10 | 3.00 | X |
| *Jaculus jaculus* | 92.80 | 3.30 | X |
| *Meriones unguiculatus* | 92.80 | 3.00 | X |
| *Mesocricetus auratus* | 90.50 | 3.50 | X |
| *Microtus ochrogaster* | 95.20 | 3.50 | X |
| *Mus musculus* | 95.20 | 2.40 | X |
| *Nannospalax galili* | 59.10 | 15.60 | |
| *Neotoma lepida* | 78.00 | 10.60 | X |
| *Onychomys torridus* | 63.40 | 11.70 | |
| *Peromyscus attwateri* | 66.40 | 11.00 | |
| *Peromyscus aztecus* | 61.70 | 12.70 | |
| *Peromyscus eremicus* | 92.90 | 3.70 | X |
| *Peromyscus leucopus* | 94.50 | 3.50 | X |
| *Peromyscus maniculatus* | 95.30 | 2.40 | X |
| *Peromyscus melanophrys* | 55.10 | 16.00 | |
| *Peromyscus nudipes* | 12.30 | 12.30 | |
| *Peromyscus polionotus* | 95.40 | 2.20 | X |
| *Phodopus sungorus* | 12.70 | 52.20 | |
| *Rattus norvegicus* | 91.60 | 3.80 | X |
| *Rhizomys pruinosus* | 14.50 | 53.40 | |
| *Sigmodon hispidus* | 86.6 | 4.8 | X |

**Supplementary Table 2.** Proportion of the genome covered by different kinds of repetitive elements. 'Unclass.' stands for 'Unclassified', 'Low compl.' stands for 'Low complexity'.

| Species | SINEs | LINEs | LTRs | DNA Elements | Unclass. | Satellites | Simple repeats | Low compl. | Total masked |
|---|---|---|---|---|---|---|---|---|---|
| *Peromyscus attwateri* | 9.36 | 11.53 | 10.74 | 1.39 | 0.5 | 0.08 | 2.17 | 0.29 | 36.16 |
| *Peromyscus aztecus* | 9.25 | 11.06 | 10.44 | 1.38 | 0.48 | 0.08 | 2.08 | 0.29 | 35.14 |
| ***Peromyscus eremicus*** | **9.22** | **10.54** | **10.51** | **1.45** | **0.46** | **0.08** | **2.07** | **0.28** | **34.68** |
| *Peromyscus leucopus* | 9.42 | 11.44 | 11.28 | 1.41 | 0.47 | 0.09 | 2.24 | 0.31 | 36.73 |
| *Peromyscus maniculatus* | 9.04 | 10.44 | 10.33 | 1.38 | 0.47 | 0.07 | 2.11 | 0.28 | 34.19 |
| *Peromyscus melanophrys* | 9.31 | 11.31 | 10.72 | 1.38 | 0.49 | 0.09 | 2.29 | 0.29 | 35.95 |
| *Peromyscus nudipes* | 9.3 | 11.37 | 10.73 | 1.39 | 0.49 | 0.08 | 2.05 | 0.28 | 35.78 |
| *Peromyscus polionotus* | 8.17 | 9.55 | 9.42 | 1.3 | 0.43 | 0.06 | 1.91 | 0.26 | 31.16 |

**Supplementary Table 3.** List of genes associated with selective sweeps in *Sweepfinder2*. 'chr' stands for 'chromosome'. Each gene was associated with its putative function using the UniProt database using *Mus musculus* as reference when available, otherwise the closest species.

| chr | start | end | Gene id | putative function |
|---|---|---|---|---|
| 1 | 5,340,876 | 5,346,674 | PRAMEF8 | regulator of apoptosis/differentiation |
| 1 | 5,449,752 | 5,460,807 | Vmn2r116 | response to pheromone |
| 1 | 10,351,947 | 10,360,112 | Vmn2r116 | response to pheromone |
| 1 | 10,361,197 | 10,361,532 | PUF | N/A |
| 1 | 16,516,689 | 16,519,154 | Usp29 | Ubl conjugation pathway |
| 1 | 16,600,642 | 16,630,107 | Usp29 | Ubl conjugation pathway |
| 1 | 16,762,270 | 16,764,639 | Usp29 | Ubl conjugation pathway |
| 1 | 16,897,493 | 16,899,184 | ZNF551 | zinc-finger, regulation of transcription |
| 1 | 16,902,915 | 16,903,847 | VN1R1 | response to pheromone |
| 1 | 16,906,907 | 16,907,839 | VN1R1 | response to pheromone |
| 1 | 16,925,094 | 16,931,066 | ZNF773 | zinc-finger, regulation of transcription |
| 1 | 16,960,660 | 16,964,629 | PUF | N/A |
| 1 | 17,003,809 | 17,007,330 | ZNF773 | zinc-finger, regulation of transcription |
| 1 | 17,024,120 | 17,024,866 | VN1R1 | response to pheromone |
| 1 | 18,248,730 | 18,249,650 | VN1R2 | response to pheromone |
| 1 | 18,362,159 | 18,362,503 | VN1R4 | response to pheromone |
| 1 | 18,745,185 | 18,745,676 | VN1R4 | response to pheromone |
| 1 | 18,952,112 | 18,952,459 | RPL23A | ribosomal protein |
| 1 | 19,015,363 | 19,016,073 | Gm5592 | unknown |
| 1 | 19,151,013 | 19,151,963 | VN1R1 | response to pheromone |
| 1 | 20,917,147 | 20,934,912 | Similar to NAL212_1203 | alcohol sulfotransferase |
| 1 | 20,971,673 | 20,984,405 | Sult2a1 | lipid metabolism |
| 1 | 21,000,532 | 21,015,370 | Similar to NAL212_1203 | alcohol sulfotransferase |
| 1 | 21,034,561 | 21,053,565 | Sult2a1 | lipid metabolism |
| 1 | 21,098,100 | 21,111,214 | Probable | N/A |
| 1 | 21,202,187 | 21,206,810 | Sult2a1 | lipid metabolism |
| 1 | 29,594,123 | 29,601,907 | Rbmy1b | mRNA splicing/spermatogenesis |
| 1 | 29,738,254 | 29,740,926 | PHF8 | chromatin regulator |
| 1 | 42,440,219 | 42,471,011 | Ush1c | hearing |
| 1 | 42,665,071 | 42,666,762 | Myod1 | muscle differentiation |
| 1 | 43,053,590 | 43,054,438 | Mrgprb2 | airway inflammation |
| 1 | 43,159,412 | 43,185,178 | Mrgprx2 | sensation or modulation of pain |
| 1 | 47,459,525 | 47,459,743 | GAS2 | growth arrest |
| 1 | 47,728,447 | 47,728,527 | PUF | N/A |
| 1 | 48,135,241 | 48,135,342 | PUF | N/A |

| | | | | |
|---|---|---|---|---|
| 1 | 48,613,009 | 48,613,146 | PUF | N/A |
| 1 | 54,742,203 | 54,743,414 | Zcchc3 | antiviral defense |
| 1 | 56,372,665 | 56,373,642 | Ndn | growth regulation |
| 1 | 100,574,169 | 100,579,965 | Dchs1 | morphogenesis |
| 1 | 100,662,592 | 100,669,716 | GVINP1 | GTP binding |
| 1 | 107,063,033 | 107,080,498 | PARVA | angiogenesis |
| 1 | 107,098,264 | 107,122,880 | Parva | angiogenesis |
| 1 | 107,413,647 | 107,452,328 | Tead1 | organ size control |
| 1 | 152,526,799 | 152,527,707 | OR9I1 | olfactory receptor |
| 1 | 152,626,688 | 152,628,403 | OR9I1 | olfactory receptor |
| 2 | 133,831 | 135,126 | TUBA4A | neuron migration |
| 2 | 20,784,083 | 20,814,269 | Mmp16 | bone development |
| 2 | 21,512,432 | 21,513,535 | HNRNPA3 | ribosomal protein |
| 2 | 45,350,465 | 45,353,607 | Nkain3 | regulation of sodium ion transport |
| 2 | 45,949,971 | 45,953,751 | Ndufaf4 | regulation of cell proliferation |
| 2 | 90,093,164 | 90,093,247 | PUF | N/A |
| 2 | 90,953,291 | 90,953,422 | PUF | N/A |
| 2 | 128,658,791 | 128,665,061 | TMEM61 | transmembrane protein |
| 2 | 128,820,350 | 128,851,896 | Dhcr24 | cholesterol metabolism |
| 3 | 70,233,195 | 70,233,618 | PUF | N/A |
| 3 | 70,883,642 | 70,905,640 | Avl9 | cell migration |
| 3 | 81,769,388 | 81,769,666 | PUF | N/A |
| 3 | 81,891,465 | 81,922,817 | Immt | mitochondrial homeostasis |
| 3 | 85,268,392 | 85,287,444 | Cntnap2 | neuronal development |
| 3 | 85,782,046 | 85,782,324 | Tox3 | apoptosis |
| 3 | 152,775,925 | 152,779,387 | Klrk1 | adaptive immunity |
| 3 | 153,062,333 | 153,063,244 | Tas2r117 | sensory perception of bitter taste |
| 4 | 159533 | 194215 | Ano3 | Lipid transport/detection of stimulus |
| 4 | 460558 | 466670 | ANKRD18A | protein-protein interactions |
| 4 | 82041414 | 82043701 | HPCAL4 | signal transduction - vision |
| 4 | 82330214 | 82354893 | Ptpa | protein folding |
| 4 | 90966365 | 90969369 | Abo | protein glycosylation |
| 4 | 92905910 | 92933519 | Lrp1b | in-utero embryonic development |
| 4 | 94065463 | 94068752 | Prss2 | digestion |
| 4 | 94927354 | 94927566 | PUF | N/A |
| 4 | 134173046 | 134173207 | PUF | N/A |
| 4 | 134566770 | 134616323 | Fsip2 | spermatogenesis |
| 5 | 55,042,327 | 55045495 | Dbndd1 | unknown |
| 5 | 55,331,567 | 55331947 | Hist1h2bf | Nucleosome formation |
| 5 | 78,686,173 | 78697206 | FAM107B | sensory perception of sound |

| | | | | |
|---|---|---|---|---|
| 5 | 79,580,816 | 79629110 | Frmd4a | establishment of epithelial cell polarity |
| 6 | 43,833,270 | 43,833,590 | Erh | transcripton and translation regulation |
| 6 | 43,846,724 | 43,847,041 | Erh | transcripton and translation regulation |
| 6 | 44,159,842 | 44,160,057 | PUF | N/A |
| 6 | 44,840,465 | 44,871,806 | ANKRD50 | protein transport |
| 6 | 55,098,083 | 55,098,235 | PUF | N/A |
| 6 | 55,760,127 | 55,760,441 | Erh | transcripton and translation regulation |
| 6 | 59,234,415 | 59,235,671 | Foxo1 | Transcription factor that is the main target of insulin signaling and regulates metabolic homeostasis in response to oxidative stress |
| 6 | 59,670,893 | 59,677,189 | Stoml3 | signal transduction - mechanoreception |
| 6 | 66,250,902 | 66,287,844 | AADACL2 | carboxylic ester hydrolase activity |
| 6 | 66,326,736 | 66,352,794 | AADACL2 | carboxylic ester hydrolase activity |
| 6 | 77,262,270 | 77,262,470 | PUF | N/A |
| 6 | 77,453,371 | 77,453,553 | TIMM23 | protein transport |
| 6 | 82,249,032 | 82,249,241 | RPL38 | ribosomal protein |
| 6 | 82,406,351 | 82,438,826 | Fstl5 | cell differentiation |
| 6 | 83,069,401 | 83,069,598 | PUF | N/A |
| 6 | 125,475,940 | 125,476,437 | OR52R1 | olfactory receptor |
| 6 | 125,525,939 | 125,549,901 | Clca4a | chloride transport |
| 7 | 15,210,037 | 15,224,726 | Vmn2r116 | response to pheromones |
| 7 | 15,257,866 | 15,327,971 | OR7G2 | olfactory receptor |
| 7 | 15,577,617 | 15,594,192 | OR7G1 | olfactory receptor |
| 7 | 15,613,673 | 15,618,557 | OR7G2 | olfactory receptor |
| 7 | 87,688,179 | 87,718,292 | SYNCRIP | RNA-binding |
| 7 | 88,037,337 | 88,037,477 | CCT3 | protein chaperone |
| 7 | 125,366,966 | 125,381,012 | Ccr2 | Inflammatory response |
| 8 | 25,405 | 42,722 | Meiob | meiosis |
| 8 | 101,476 | 101,607 | PUF | N/A |
| 8 | 3,625,875 | 3,646,917 | Abcc1 | inflammatory response |
| 8 | 3,660,678 | 3,705,621 | Abcc1 | inflammatory response |
| 8 | 37,085,591 | 37,086,481 | PUF | N/A |
| 8 | 37,135,633 | 37,148,801 | CLK4 | regulation of RNA splicing |
| 8 | 37,528,225 | 37,532,843 | HNRNPAB | Transcription regulation |
| 8 | 37,600,330 | 37,608,225 | Rmnd5b | Ubl conjugation pathway |
| 8 | 37,629,246 | 37,635,416 | N4bp3 | Neurogenesis |
| 8 | 37,672,929 | 37,676,601 | TRAPPC2 | skeletal system development |
| 8 | 45,907,237 | 45,910,943 | Wnt9a | embryonic bone development |
| 8 | 45,960,544 | 45,962,754 | Prss38 | protease |
| 8 | 53,487,636 | 53,488,034 | DNAH9 | pancreas development |
| 8 | 53,594,370 | 53,635,391 | DNAH9 | pancreas development |

| | | | | |
|---|---|---|---|---|
| 9 | 21349153 | 21366743 | Gdf10 | adipogenesis and osteogenesis |
| 9 | 22022099 | 22022686 | PUF | N/A |
| 9 | 29461042 | 29461110 | PUF | N/A |
| 9 | 29646610 | 29646645 | PUF | N/A |
| 9 | 29868982 | 29895666 | Msmo1 | lipid metabolism |
| 9 | 30061380 | 30067261 | Ppif | apoptosis, necrosis |
| 9 | 32142451 | 32142486 | PUF | N/A |
| 9 | 32160796 | 32162383 | PUF | N/A |
| 9 | 45652331 | 45652681 | TRDV1 | immune response |
| 9 | 45663675 | 45664025 | TRAV14DV4 | immune response |
| 9 | 70079245 | 70080354 | Spert | spermatid formation |
| 9 | 70238094 | 70238360 | RPL30 | defence response to bacteria |
| 10 | 48473687 | 48474013 | PUF | N/A |
| 10 | 48655763 | 48655861 | PUF | N/A |
| 10 | 103846797 | 103854163 | Chac2 | glutathione catabolic process |
| 11 | 8273550 | 8273789 | PUF | N/A |
| 11 | 10252681 | 10252869 | rpl15 | ribosomal protein |
| 11 | 10482956 | 10483039 | PUF | N/A |
| 11 | 12505603 | 12553690 | CNTNAP5 | development and functioning of nervous system |
| 11 | 13333604 | 13333663 | PUF | N/A |
| 11 | 13508953 | 13509012 | PUF | N/A |
| 11 | 13927456 | 13927755 | PUF | N/A |
| 11 | 14245488 | 14245961 | HRAS | GTPase - many processes |
| 11 | 15023224 | 15023451 | Gm5592 | uncharacterized protein' |
| 11 | 15728361 | 15728814 | PUF | N/A |
| 11 | 17132226 | 17132390 | PUF | N/A |
| 11 | 17200894 | 17200926 | PUF | N/A |
| 11 | 17287535 | 17288948 | RPAP3 | regulator of protein complex formation |
| 11 | 17511597 | 17511871 | Rpl7a | RNA binding |
| 11 | 17575392 | 17588436 | Snapc1 | transcription regulation |
| 11 | 17793024 | 17793776 | Hnrnpa1 | mRNA processing |
| 11 | 18957115 | 18961723 | TSN | DNA- and RNA-binding |
| 11 | 65578533 | 65578953 | RPL32 | ribosomal protein |
| 11 | 66361669 | 66367491 | Gpr161 | developmental protein |
| 11 | 66386947 | 66389163 | PUF | N/A |
| 11 | 66572541 | 66639559 | Dcaf6 | transcription regulation |
| 12 | 13933193 | 13937526 | Rtp2 | sensory perception of bitter taste |
| 12 | 14153756 | 14162594 | Rtp4 | sensory perception of bitter taste |
| 12 | 76417361 | 76449259 | DNM1L | biological rhythms |
| 12 | 76930895 | 76931131 | Olfr19 | sensory perception of smell |

| | | | | |
|---|---|---|---|---|
| 12 | 79125272 | 79152393 | Senp7 | ubiquitin-like modifier processing |
| 12 | 79404693 | 79405106 | SUMO1 | stress response (including cellular repsonse to heat) |
| 13 | 5047275 | 5047853 | ARL4C | GTP-binding (cholesterol secretion pathway) |
| 13 | 5545804 | 5566717 | Sh3bp4 | endocytosis |
| 13 | 5757375 | 5757512 | PUF | N/A |
| 13 | 6158478 | 6210352 | Agap1 | protein transport |
| 14 | 38187366 | 38211373 | Ig heavy chain | adaptive immunity |
| 14 | 38273092 | 38299292 | Igh-VJ558 | adaptive immunity |
| 15 | 10090784 | 10098139 | PUF | N/A |
| 15 | 10314805 | 10315284 | HMGN2 | regulation of transcription |
| 15 | 11619868 | 11620101 | PUF | N/A |
| 15 | 11992254 | 11992352 | PUF | N/A |
| 15 | 24858214 | 24858354 | PUF | N/A |
| 15 | 26128400 | 26128492 | RPL21 | transcription regulation |
| 16 | 32021644 | 32040753 | SLC18B1 | solute carrier |
| 16 | 32105692 | 32121221 | Vnn1 | inflammatory response |
| 17 | 27823141 | 27837356 | Fam149a | function unknown |
| 17 | 28006571 | 28027817 | Cyp4v2 | lipid metabolism |
| 17 | 62570441 | 62624946 | Ikbkb | NF-kappa-B signaling pathway, activated by cellular stress (including pathogens) |
| 17 | 62703490 | 62716746 | Ap3m2 | protein transport |
| 18 | 3410632 | 3445879 | Aldh1l2 | oxidoreductase activity |
| 18 | 4024104 | 4024925 | Chst11 | cartilage development |
| 19 | 3147121 | 3195632 | Npc1 | Cholesterol metabolism |
| 19 | 3470976 | 3529638 | Lama3 | regulation of embryomnic development |
| 20 | 17867368 | 17870548 | Fbxl6 | Ubl conjugation pathway |
| 20 | 17913321 | 17920126 | Adck5 | function unknown |
| 21 | 18076629 | 18095946 | Mog | regulation of membrane potential |
| 21 | 18224976 | 18225578 | Znrd1-as | may be involved in male sterility |
| 21 | 62653400 | 62669212 | Mep1a | zinc ion binding |
| 21 | 63159390 | 63161863 | GSTM4 | glutathione metabolic processing |
| 22 | 19104429 | 19105207 | PUF | N/A |
| 22 | 19744798 | 19751653 | Ntsr2 | regulation of membrane potential |
| 22 | 48879348 | 48919777 | Vmn2r116 | response to pheromones |
| 22 | 48925556 | 48925950 | RPS2 | ribosomal protein |
| 22 | 52936806 | 52938793 | Npm1 | DNA- and RNA-binding |
| 22 | 53103853 | 53104173 | Vmn2r116 | response to pheromones |
| 22 | 54093646 | 54094167 | CBX3 | chromatin organization, biological rhythms |
| 22 | 54881239 | 54938624 | Adgre1 | adaptive immune response |
| 22 | 58549681 | 58553574 | Plpp2 | phospholipid methabolism |

| | | | | |
|---|---|---|---|---|
| 22 | 58669486 | 58669602 | PUF | N/A |
| 23 | 36650892 | 36671552 | Fam20c | biomineralization (bone formation) |
| 23 | 36827756 | 36846546 | Vmn2r116 | response to pheromones |

**Supplementary Table 4.** Enriched GO terms for biological processes associated with selective sweeps reduced by semantic similarity in *REVIGO*.

| GO term ID | Description (biological processes) | log10 p-value |
|---|---|---|
| GO:0061024 | membrane organization | -3.6517 |
| GO:0043603 | cellular amide metabolic process | -3.5622 |
| GO:0006412 | translation | -3.4962 |
| GO:0042255 | ribosome assembly | -3.2636 |
| GO:0051346 | negative regulation of hydrolase activity | -2.8153 |
| GO:0001580 | detection of chemical stimulus involved in sensory perception of bitter taste | -2.5884 |
| GO:0009987 | cellular process | -2.5003 |
| GO:0071243 | cellular response to arsenic-containing substance | -2.4353 |
| GO:0060548 | negative regulation of cell death | -2.4023 |
| GO:0042219 | cellular modified amino acid catabolic process | -2.295 |
| GO:0033036 | macromolecule localization | -2.2916 |
| GO:0035329 | hippo signaling | -2.2534 |
| GO:0070266 | necroptotic process | -2.2534 |
| GO:0098542 | defense response to other organism | -2.2299 |
| GO:0050909 | sensory perception of taste | -2.1494 |
| GO:0043484 | regulation of RNA splicing | -2.1325 |
| GO:0008037 | cell recognition | -2.1267 |
| GO:0048562 | embryonic organ morphogenesis | -2.1203 |
| GO:0071359 | cellular response to dsRNA | -2.104 |
| GO:0032729 | positive regulation of interferon-gamma production | -2.0857 |
| GO:0046685 | response to arsenic-containing substance | -2.0711 |
| GO:0006897 | endocytosis | -2.065 |
| GO:0036342 | post-anal tail morphogenesis | -2.0391 |
| GO:0098657 | import into cell | -2.0362 |
| GO:1902629 | regulation of mRNA stability involved in cellular response to UV | -1.9706 |
| GO:0007518 | myoblast fate determination | -1.9706 |
| GO:2000818 | negative regulation of myoblast proliferation | -1.9706 |
| GO:0036179 | osteoclast maturation | -1.9706 |
| GO:0071716 | leukotriene transport | -1.9706 |
| GO:1905380 | regulation of snRNA transcription from RNA polymerase II promoter | -1.9706 |
| GO:0035705 | T-helper 17 cell chemotaxis | -1.9706 |
| GO:0010849 | regulation of proton-transporting ATPase activity rotational mechanism | -1.9706 |
| GO:0002253 | activation of immune response | -1.8962 |
| GO:0070265 | necrotic cell death | -1.8962 |
| GO:0030574 | collagen catabolic process | -1.8447 |

| | | |
|---|---|---|
| GO:0051205 | protein insertion into membrane | -1.821 |
| GO:0070887 | cellular response to chemical stimulus | -1.8013 |
| GO:0007585 | respiratory gaseous exchange | -1.7986 |
| GO:0046686 | response to cadmium ion | -1.7986 |
| GO:1904106 | protein localization to microvillus | -1.7959 |
| GO:2000473 | positive regulation of hematopoietic stem cell migration | -1.7959 |
| GO:0010430 | fatty acid omega-oxidation | -1.7959 |
| GO:0033037 | polysaccharide localization | -1.7959 |
| GO:0001544 | initiation of primordial ovarian follicle growth | -1.7959 |
| GO:1900063 | regulation of peroxisome organization | -1.7959 |
| GO:0035672 | oligopeptide transmembrane transport | -1.7959 |
| GO:0035574 | histone H4-K20 demethylation | -1.7959 |
| GO:0061188 | negative regulation of chromatin silencing at rDNA | -1.7959 |
| GO:1904578 | response to thapsigargin | -1.7959 |
| GO:0090149 | mitochondrial membrane fission | -1.7959 |
| GO:2000458 | regulation of astrocyte chemotaxis | -1.7959 |
| GO:0090265 | positive regulation of immune complex clearance by monocytes and macrophages | -1.7959 |
| GO:0015939 | pantothenate metabolic process | -1.7959 |
| GO:1904579 | cellular response to thapsigargin | -1.7959 |
| GO:0099039 | sphingolipid translocation | -1.7959 |
| GO:0001501 | skeletal system development | -1.767 |
| GO:0010467 | gene expression | -1.7282 |
| GO:0043503 | skeletal muscle fiber adaptation | -1.6737 |
| GO:1902349 | response to chloroquine | -1.6737 |
| GO:0090204 | protein localization to nuclear pore | -1.6737 |
| GO:0060699 | regulation of endoribonuclease activity | -1.6737 |
| GO:0019538 | protein metabolic process | -1.6536 |
| GO:0045740 | positive regulation of DNA replication | -1.6345 |
| GO:0001738 | morphogenesis of a polarized epithelium | -1.58 |
| GO:0018916 | nitrobenzene metabolic process | -1.5768 |
| GO:0071895 | odontoblast differentiation | -1.5768 |
| GO:0072137 | condensed mesenchymal cell proliferation | -1.5768 |
| GO:0097350 | neutrophil clearance | -1.5768 |
| GO:1903347 | negative regulation of bicellular tight junction assembly | -1.5768 |
| GO:0003273 | cell migration involved in endocardial cushion formation | -1.5768 |
| GO:0022613 | ribonucleoprotein complex biogenesis | -1.5735 |
| GO:0007166 | cell surface receptor signaling pathway | -1.5686 |
| GO:0051438 | regulation of ubiquitin-protein transferase activity | -1.5302 |
| GO:0016043 | cellular component organization | -1.5258 |

| | | |
|---|---|---|
| GO:0001503 | ossification | -1.5186 |
| GO:0006952 | defense response | -1.5045 |
| GO:0046618 | drug export | -1.4989 |
| GO:0008038 | neuron recognition | -1.4989 |
| GO:1904970 | brush border assembly | -1.4989 |
| GO:0003192 | mitral valve formation | -1.4989 |
| GO:0071205 | protein localization to juxtaparanode region of axon | -1.4989 |
| GO:0097187 | dentinogenesis | -1.4989 |
| GO:0009597 | detection of virus | -1.4989 |
| GO:0071840 | cellular component organization or biogenesis | -1.4962 |
| GO:0071826 | ribonucleoprotein complex subunit organization | -1.4949 |
| GO:0051179 | localization | -1.4737 |
| GO:0097035 | regulation of membrane lipid distribution | -1.4698 |
| GO:0051606 | detection of stimulus | -1.4461 |
| GO:0006807 | nitrogen compound metabolic process | -1.4449 |
| GO:0045925 | positive regulation of female receptivity | -1.433 |
| GO:0055093 | response to hyperoxia | -1.433 |
| GO:0036466 | synaptic vesicle recycling via endosome | -1.433 |
| GO:1904751 | positive regulation of protein localization to nucleolus | -1.433 |
| GO:0002544 | chronic inflammatory response | -1.433 |
| GO:0008202 | steroid metabolic process | -1.4295 |
| GO:0090303 | positive regulation of wound healing | -1.4123 |
| GO:0034645 | cellular macromolecule biosynthetic process | -1.4056 |
| GO:0042221 | response to chemical | -1.382 |
| GO:0032532 | regulation of microvillus length | -1.3768 |
| GO:0030046 | parallel actin filament bundle assembly | -1.3768 |
| GO:1903608 | protein localization to cytoplasmic stress granule | -1.3768 |
| GO:0042297 | vocal learning | -1.3768 |
| GO:0007569 | cell aging | -1.3605 |
| GO:0009605 | response to external stimulus | -1.3458 |
| GO:0010506 | regulation of autophagy | -1.3363 |
| GO:0010501 | RNA secondary structure unwinding | -1.3261 |
| GO:0090324 | negative regulation of oxidative phosphorylation | -1.3261 |
| GO:0007144 | female meiosis I | -1.3261 |
| GO:1901620 | regulation of smoothened signaling pathway involved in dorsal/ventral neural tube patterning | -1.3261 |
| GO:1901564 | organonitrogen compound metabolic process | -1.3224 |
| GO:1901698 | response to nitrogen compound | -1.3206 |
| GO:1903311 | regulation of mRNA metabolic process | -1.3152 |

**Supplementary Table 5.** Enriched GO terms for molecular function associated with selective sweeps reduced by semantic similarity in *REVIGO*.

| GO term ID | description (molecular functions) | log10 p-value |
|---|---|---|
| GO:0003735 | structural constituent of ribosome | -4.3686 |
| GO:0005488 | binding | -3.2865 |
| GO:0031849 | olfactory receptor binding | -3.2321 |
| GO:0003729 | mRNA binding | -3.2248 |
| GO:0003723 | RNA binding | -2.7496 |
| GO:0001221 | transcription cofactor binding | -2.5406 |
| GO:0043024 | ribosomal small subunit binding | -2.4353 |
| GO:0097159 | organic cyclic compound binding | -2.3468 |
| GO:0022884 | macromolecule transmembrane transporter activity | -2.295 |
| GO:1901363 | heterocyclic compound binding | -2.2321 |
| GO:0005198 | structural molecule activity | -2.2226 |
| GO:0008143 | poly(A) binding | -2.0083 |
| GO:0000246 | delta24(24-1) sterol reductase activity | -1.9706 |
| GO:1990595 | mast cell secretagogue receptor activity | -1.9706 |
| GO:0035715 | chemokine (C-C motif) ligand 2 binding | -1.9706 |
| GO:0051433 | BH2 domain binding | -1.9706 |
| GO:0004381 | fucosylgalactoside 3-alpha-galactosyltransferase activity | -1.9706 |
| GO:0043273 | CTPase activity | -1.9706 |
| GO:0034634 | glutathione transmembrane transporter activity | -1.9706 |
| GO:0003823 | antigen binding | -1.9318 |
| GO:0005515 | protein binding | -1.8239 |
| GO:0008160 | protein tyrosine phosphatase activator activity | -1.7959 |
| GO:0032029 | myosin tail binding | -1.7959 |
| GO:0035575 | histone demethylase activity (H4-K20 specific) | -1.7959 |
| GO:0017159 | pantetheine hydrolase activity | -1.7959 |
| GO:0032394 | MHC class Ib receptor activity | -1.7959 |
| GO:0050659 | N-acetylgalactosamine 4-sulfate 6-O-sulfotransferase activity | -1.7959 |
| GO:0016492 | G-protein coupled neurotensin receptor activity | -1.7959 |
| GO:0000254 | C-4 methylsterol oxidase activity | -1.7959 |
| GO:0016155 | formyltetrahydrofolate dehydrogenase activity | -1.7959 |
| GO:1904841 | TORC2 complex binding | -1.7959 |
| GO:0034040 | lipid-transporting ATPase activity | -1.7959 |
| GO:0051082 | unfolded protein binding | -1.7799 |
| GO:0003755 | peptidyl-prolyl cis-trans isomerase activity | -1.7545 |
| GO:0051721 | protein phosphatase 2A binding | -1.7122 |
| GO:0004930 | G-protein coupled receptor activity | -1.6861 |

| | | |
|---|---|---|
| GO:0070139 | SUMO-specific endopeptidase activity | -1.6737 |
| GO:0008384 | IkappaB kinase activity | -1.6737 |
| GO:0003839 | gamma-glutamylcyclotransferase activity | -1.6737 |
| GO:0051033 | RNA transmembrane transporter activity | -1.6737 |
| GO:0003924 | GTPase activity | -1.6596 |
| GO:0042887 | amide transmembrane transporter activity | -1.6345 |
| GO:0034235 | GPI anchor binding | -1.5768 |
| GO:0034056 | estrogen response element binding | -1.5768 |
| GO:0001134 | transcription factor activity, transcription factor recruiting | -1.4989 |
| GO:0008310 | single-stranded DNA 3'-5' exodeoxyribonuclease activity | -1.4989 |
| GO:0044388 | small protein activating enzyme binding | -1.4989 |
| GO:0003676 | nucleic acid binding | -1.4789 |
| GO:0015562 | efflux transmembrane transporter activity | -1.433 |
| GO:0016842 | amidine-lyase activity | -1.3768 |
| GO:0050656 | 3'-phosphoadenosine 5'-phosphosulfate binding | -1.3768 |
| GO:0030145 | manganese ion binding | -1.3726 |
| GO:0005525 | GTP binding | -1.3516 |
| GO:0016742 | hydroxymethyl-, formyl- and related transferase activity | -1.3261 |
| GO:0005319 | lipid transporter activity | -1.3224 |
| GO:0005509 | calcium ion binding | -1.3197 |

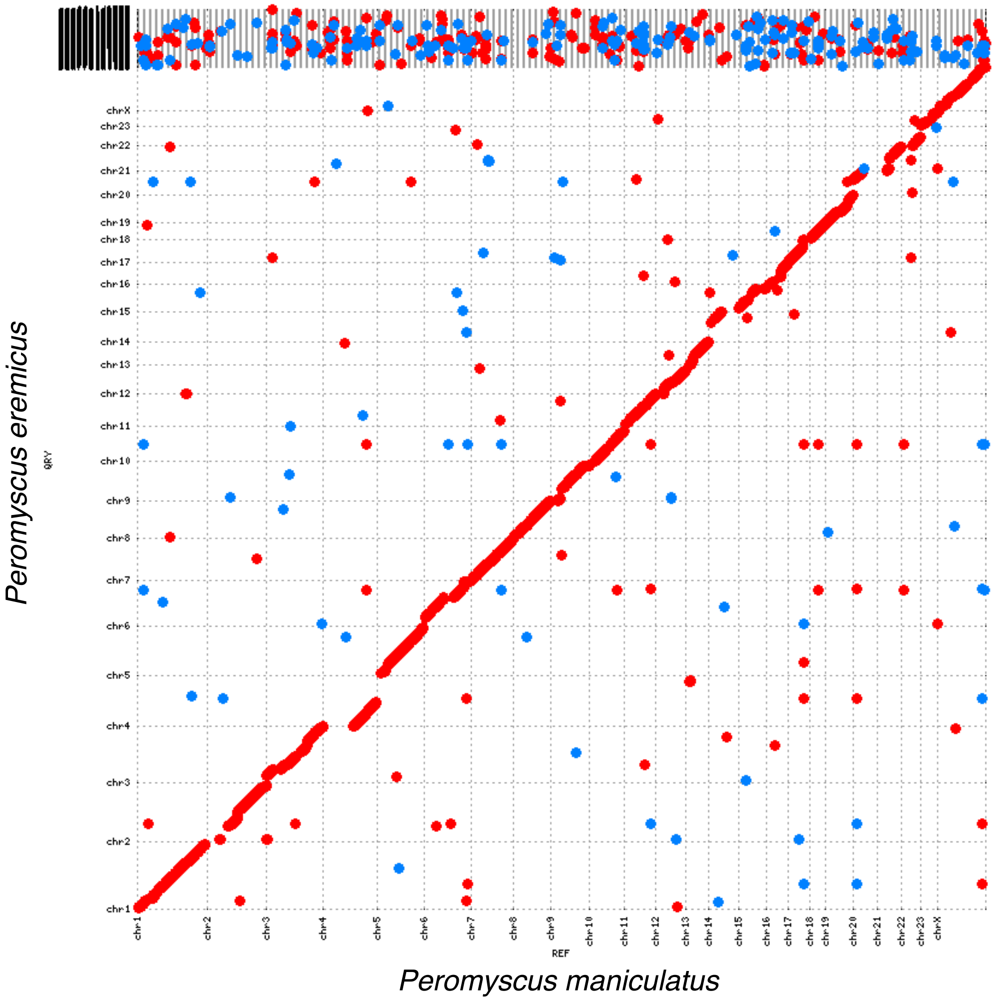


**Supplementary Figure 1.** Dotplot showing syntenic relationships between chromosomes of *Peromyscus eremicus* on the y axis and *Peromyscus maniculatus* on the x axis.

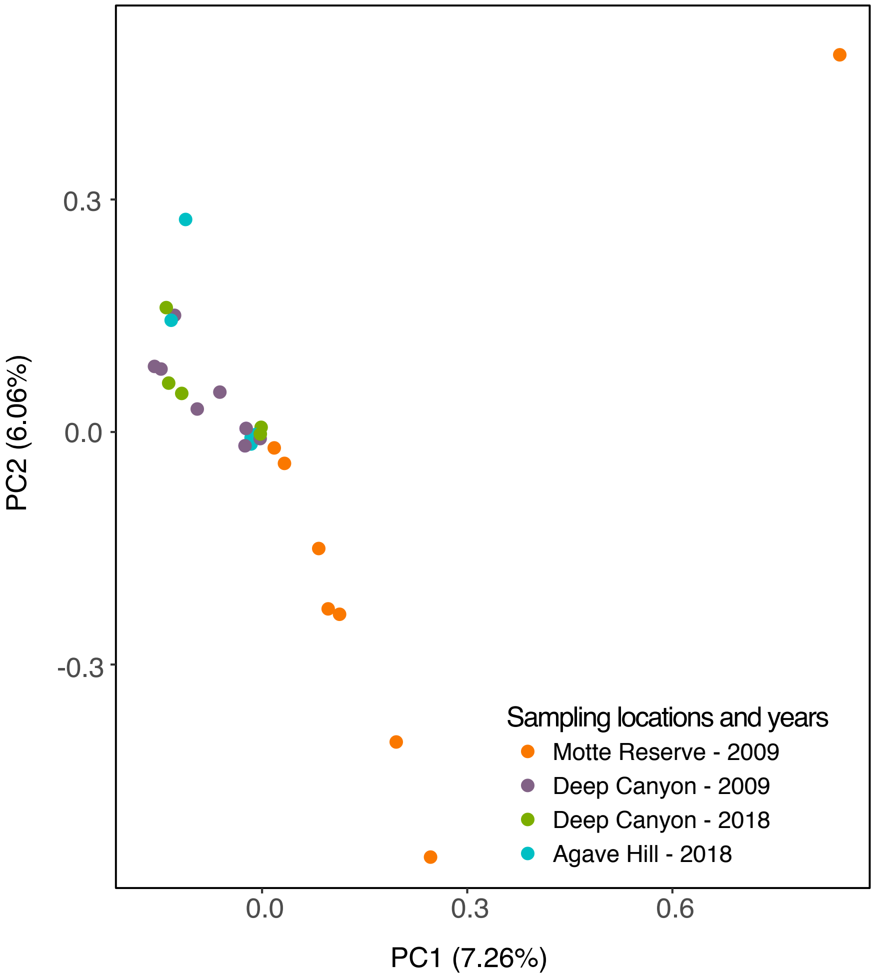


**Supplementary Figure 2.** PCA plot showing clustering of individuals based on 43.7 million SNPs.

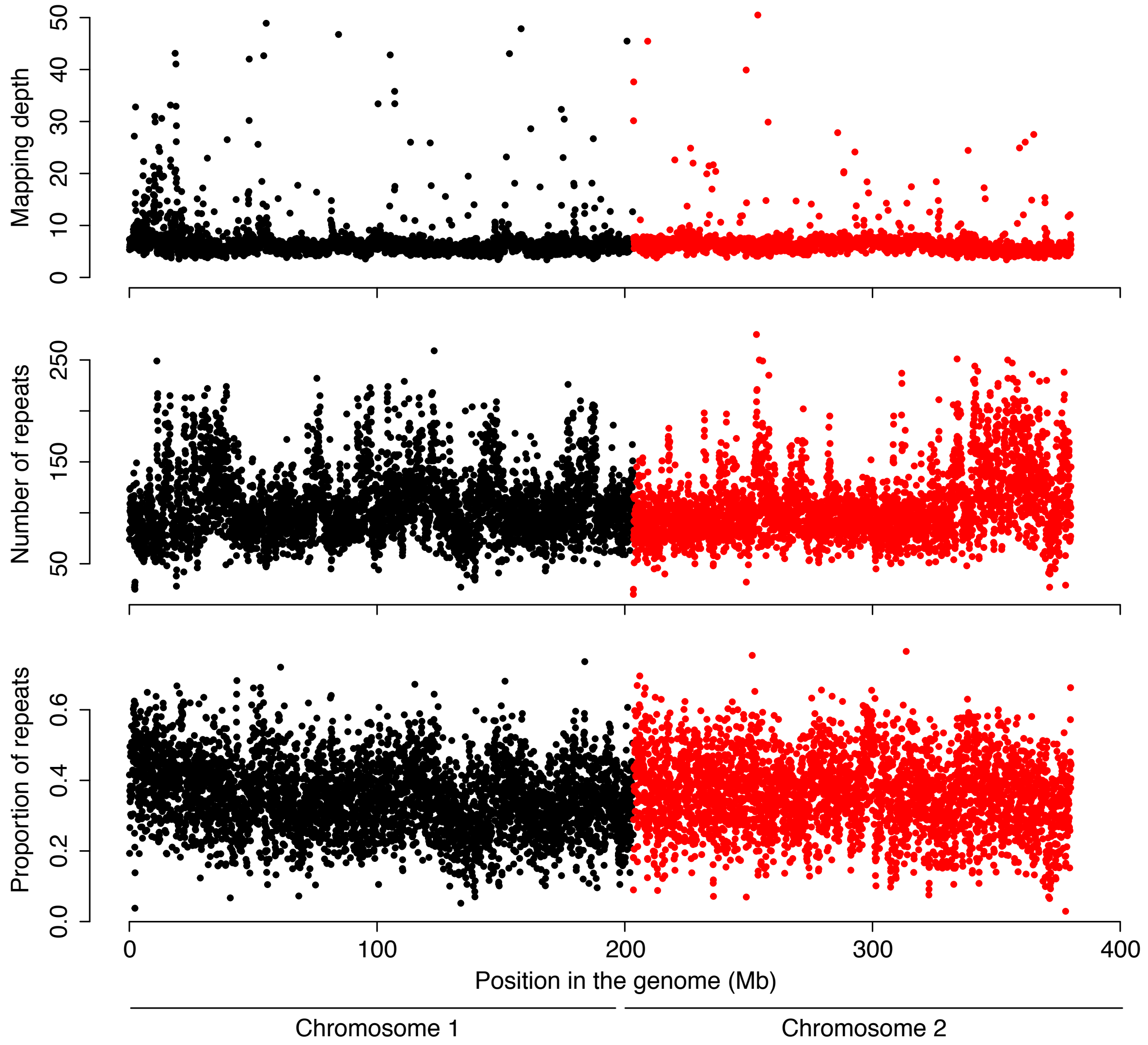


**Supplementary Figure 3.** Mapping depth, number of repeats and proportion of repeats calculated in 50 kb moving windows in chromosome 1 (in black), which shows a large peak of elevated π (Figure 1B from the main text), and chromosome 2 (in red), used as comparison for lack of π peaks. In correspondence of the π peak in the beginning on chromosome 1, mapping depth, number of repeats and proportion of repeats are also elevated when compared to the surrounding regions and/or chromosome 2.

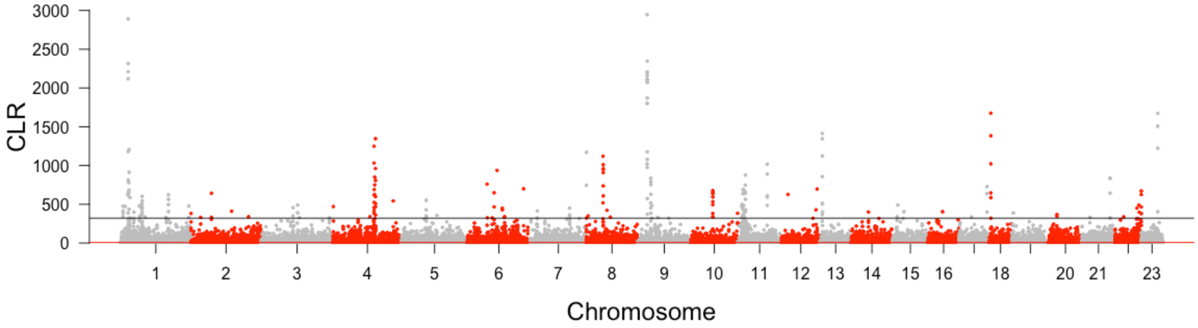


**Supplementary Figure 4.** Manhattan plot of CLR values from *Sweepfinder2*. Values over the 99.9th percentile (horizontal line) are considered significant selective sweeps.

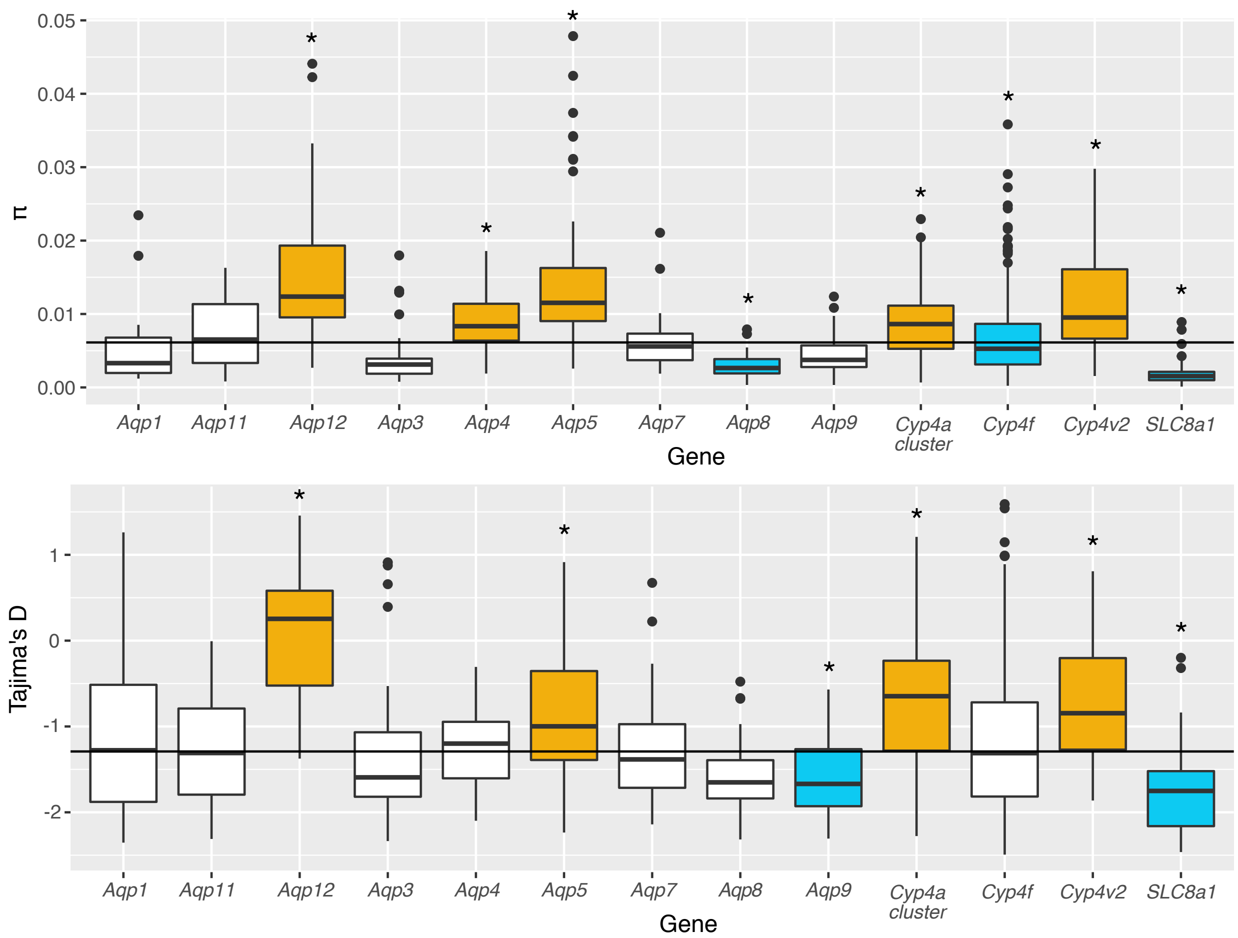


**Supplementary Figure 5.** Boxplots of π and Tajima's (upper and lower panel, respectively) calculated in 1 kb windows for each of the candidate genes identified *a priori* plus 10 kb regions flanking each direction. Stars indicate significant deviations from genome-wide means ($p < 0.05$ after Benjamini-Yekutieli correction for multiple testing), orange and blue boxes indicate respectively significantly higher and lower mean when compared to genome-wide means.
